# Ribonucleotide Reductase Regulatory Subunit M2 as a Driver of Glioblastoma TMZ-Resistance through Modulation of dNTP Production

**DOI:** 10.1101/2021.11.23.469785

**Authors:** Ella N Perrault, Jack M Shireman, Eunus S Ali, Isabelle Preddy, Peiyu Lin, Cheol Park, Luke Tomes, Andrew J Zolp, Shreya Budhiraja, Shivani Baisiwala, C. David James, Issam Ben-Sahra, Sebastian Pott, Anindita Basu, Atique U Ahmed

## Abstract

Glioblastoma (GBM) remains one of the most resistant and fatal forms of cancer. Previous studies have examined primary and recurrent GBM tumors, but it is difficult to study tumor evolution during therapy where resistance develops. To investigate this, we performed an *in vivo* single-cell RNA sequencing screen in a patient-derived xenograft (PDX) model. Primary GBM was modeled by mice treated with DMSO control, recurrent GBM was modeled by mice treated with temozolomide (TMZ), and during therapy GBM was modeled by mice euthanized after two of five TMZ treatments. Our analysis revealed the cellular population present during therapy to be distinct from primary and recurrent GBM. We found the Ribonucleotide Reductase gene family to exhibit a unique signature in our data due to an observed subunit switch to favor RRM2 during therapy. GBM cells were shown to rely on RRM2 during therapy causing RRM2-knockdown (KD) cells to be TMZ-sensitive. Using targeted metabolomics, we found RRM2-KDs to produce less dGTP and dCTP than control cells in response to TMZ (*p<0.0001*). Supplementing RRM2-KDs with deoxycytidine and deoxyguanosine rescued TMZ-sensitivity, suggesting an RRM2-driven mechanism of chemoresistance, established by regulating the production of these nucleotides. *In vivo*, tumor-bearing mice treated with the RRM2-inhibitor, Triapine, in combination with TMZ, survived longer than mice treated with TMZ alone (*p<0.01*), indicating promising clinical opportunities in targeting RRM2. Our data present a novel understanding of RRM2 activity, and its alteration during therapeutic stress as response to TMZ-induced DNA damage.

## INTRODUCTION

Terminal cancer is an intricate concoction of cellular heterogeneity and adaptation. As the disease evolves and therapeutic measures are taken, malignant cell populations adapt and resist therapies by becoming more heterogeneous. Glioblastoma multiforme (GBM), the most aggressive primary adult brain tumor, exhibits carcinogenic properties driven by cellular adaptation and therapeutic resistance, ultimately resulting in fatal recurrence^1, 2^.

Previous studies have investigated the difference between cellular subpopulations present pre-therapy and during post-tumor recurrence, but access to GBM tissue during chemotherapy is limited, which has historically hindered research^3, 4^. The intra-tumoral adaptations that arise during chemotherapy, driving therapeutic resistance and tumor recurrence, remain largely elusive. This raises the fundamental question of how the phenotypic diversity of GBM cells in relation to functional characteristics associated with therapeutic resistance is integral to remediation of this disease. To advance therapeutic approaches, it is vital to study how individual cells adapt during treatment, fueling chemoresistance and tumor recurrence in GBM.

To investigate this, our lab performed a single-cell RNA sequencing screen (scRNA-Seq). We developed an *in vivo* pipeline to profile gene expression and epigenetic shifts at the single-cell level before, during, and after therapy in a patient-derived xenograft (PDX) model of human primary GBM tumor cells. In GBM, temozolomide (TMZ) is the standard chemotherapy used^1–4^. scRNA-Seq can largely identify single cells in a heterogeneous tumor cell population and characterize the relative gene expression of every cell^5, 16^. Our GBM model utilized single-cell technology to identify clinically relevant gene expression patterns that underlie mechanisms of therapeutic resistance. Investigating genomic and cellular adaptations that arise during TMZ therapy is critical in discerning the functional and architectural foundations of therapeutic resistance, contributing to the indispensable differences that emerge between primary and recurrent GBM tumors^7, 8^. By comparing single-cell expression patterns across time points, we identified cellular populations present during TMZ-therapy distinct from primary and recurrent GBM.

The ribonucleotide reductase (RNR) family genes particularly stood out during our analysis due to their essential involvement in metabolic pathways altered during TMZ therapy. Classically, the beta subunit RRM2, or its isoform RRM2B, forms a complex with the alpha subunit RRM1 to create an RNR enzyme, which mediates dNTP production^9^. Our single-cell data revealed a compelling relationship between RNR subunits and their dependency on therapeutic progression. This study suggests that GBM cells rely on the beta subunit RRM2 during therapy to mediate TMZ-induced DNA damage. We demonstrate that RRM2 drives chemoresistance by promoting the production of two specific nucleotides – dCTP and dGTP – and the absence of RRM2 renders cells susceptible to TMZ. Further, inhibition of RRM2 activity using 3-AP Triapine, a drug used in several late-phase clinical studies^10^, significantly increases the therapeutic efficacy of TMZ *in vivo*, indicating encouraging clinical opportunities for GBM therapy. Overall, our data present a novel understanding of the role of RRM2 and how its activity is altered during therapeutic stress to counteract TMZ-induced DNA damage – a pathway that, when inhibited, shows promising clinical outcomes.

## MATERIALS & METHODS

### In Vivo Single-Cell RNA Sequencing Model

Athymic nude mice were acquired from Charles River Laboratories (nu/nu; Charles River, Skokie, IL, USA) and housed according to IACUC standards as defined by Northwestern^11^. Prior to intracranial implantation of GBM 43 PDX line all mice were confirmed anesthetized via administration of ketamine and xylazine. Tumors were placed 3 mm into the cortex of the mouse through a burr hole and incision was closed using sterile sutures. Upon recovery from anesthetic analgesics were administered and mice were observed to be bright, alert, and reactive. In total 150,000 GBM 43 cells were injected per mouse and an incubation period of 7 days was allowed in order to give the PDX cells time to establish a tumor before any therapy began.

After the conclusion of the 7-day incubation period mouse groups were divided equally based on sex to receive 2.5mg/kg TMZ or DMSO (vehicle control) given IP for a period of 5 days total. During day 3 of TMZ treatment a group of mice was sacrificed, and their tumors were biopsied and taken for single cell processing. This represents the middle time point within the study. Finally, after the 5-day dosing period was complete the mice were monitored until symptoms of the tumor became evident at which point they were sacrificed and the tumors were processed for single cell analysis. This represents the post therapy timepoint in the study.

### Single Cell Processing and DropSeq

After the mice were sacrificed due to tumor burden tumor bulk and margins were dissected out using surgical technique. Tumors were then enriched for human cells using a Miltenyi Mag bead purification kit specific to human HLA in order to separate out mouse cells that could have invaded the PDX tumors. This process was done according to the manufacturer’s instructions as supplied with the kit. After HLA purification a count of live cells was obtained, and cells were prepped to be run through the DropSeq microfluidic device.

For Drop Sequencing we followed the protocol set forth by Macosko and Basu et al. 2015^12^. Briefly cells were captured during a 15-minute droplet run on a clean microfluidic chip in an unused lane. Droplets were imaged for quality and to ensure a minimal number of doublets were present. Droplets were then broken, and library preparation was conducted for single cell sequencing.

### Bioinformatic Analysis

The bulk of the bioinformatics analysis of single cell data was done in Seurat v3.0. A custom pipeline was utilized in order to score cells on their ability to align to mouse as well as human genomes in order to bioinformatically determine if there was mouse cell invasion in our samples that was not able to be eliminated with our magnetic bead purification. The results of this pipeline produced a score that was stored in the Seurat meta-data and used to definitively subset our data into purely human cells when indicated. All single cell figures made for this manuscript can be created in base Seurat and are well annotated in the vignettes provided by the developers. In general, standard workflow for single cell processing as outlined in these vignettes was followed with data filtration based on UMI, nGene counts, and percent MitoReads. Clustering was determined by the number of principle components revealed to be significant using both elbow plots and Jackstraw methods. TSNE and UMAP were both employed for dimensional reduction of the data.

### Cell Lines and Culture

A human glioma cell line, U251, was procured from the American Type Culture Collection (Manassas, VA, USA). In order to culture the cells, Dulbecco’s Modified Eagle’s Medium (DMEM; Corning, Corning, NY, USA) –– containing 10% fetal bovine serum (FBS; Atlanta Biologicals, Lawrenceville, GA, USA) and 1% penicillin-streptomycin antibiotic mixture (Cellgro, Herndon, VA, USA; Mediatech, Herndon, VA, USA) –– was used^11^.

Patient-derived xenograft (PDX) glioma cell lines (GBM6, GBM39, and GBM43) were obtained from Dr. C. David James at Northwestern University. Cells were maintained according to protocol^11^. However, the PDX cells were cultured in DMEM composed of 1% FBS and 1% penicillin-streptomycin. A frozen stock, maintained in liquid nitrogen at -180°C in pure FBS supplemented with 10% dimethyl sulfoxide (DMSO), was utilized to replenish cells that had reached a maximum of four passages.

### In Vivo 3AP + Triapine Model

Athymic nude mice (nu/nu; Charles River, Skokie, IL, USA) were utilized in this study and housed in compliance with Institutional Animal Care and Use Committee (IACUC) requirements along with federal and state statutes^11^. Animals were housed in shoebox cages equipped with food and water and subjected to a 12-hour light and dark cycle^11^.

Intracranial implantation of glioblastoma cells was conducted according to our lab’s previously established glioblastoma mouse model^11^. The animals first received buprenex and metacam by intraperitoneal (IP) injection. They then were anesthetized from a second injection of ketamine/xylazine mixture (Henry Schein; New York, NY, USA). To confirm sedation was complete, a toe-pinch was conducted. Betadine and ethanol were applied to the scalp for sterilization, and artificial tears were applied to each eye. The skull was then exposed by creating a small incision using a scalpel, where after a ∼1mm burr hole was drilled right above the right frontal lobe. The mice were then placed in a stereotactic rig, where, over a period of one minute, 150,000 cells GBM43, GBM6, or GBM39), loaded in a Hamilton syringe, were injected 3 mm from the dura. The needle was then raised slightly and left for an additional minute in order to ensure that the cell suspension was released. The syringe was removed slowly, and the scalp was closed with sutures (Ethicon; Cincinnati, OH, USA), maintaining a consistent position of the head throughout the closing process. Animals were placed on heating pads until awake and responsive^11^.

Drug treatments were initiated one week following the implantation in the following manner: IP injections of either TMZ (2.5 mg/kg) or equimolar DMSO each day for five consecutive days. Experimental groups were formatted as follows: TMZ (3 mice), DMSO (3 mice), Triapine 20 mg/kg (3 mice), TMZ 2.5 mg/kg and Triapine 20 mg/kg every day for 5 days (7 mice), and TMZ mg/kg and Triapine 40 mg/kg every other day for 5 treatments (7 mice). Triapine injections were always administered 6 hours before TMZ in the mice that were treated for both. Throughout the treatment period, signs of tumor progression -- including weight reduction, hunching, and reduced body temperature -- were observed for and recorded. According to IACUC and Northwestern University guidelines, animals were sacrificed once it was evident that they would not survive past the next morning^11^.

### Immunofluorescence

Following euthanasia, mice were supplied with ice-cold PBS. After, their brains were frozen in cryoprotectant on dry ice, kept at -80 °C, sectioned at 8 *μ*m per section, and stained according to immunohistochemistry protocol^11^. Sections were thawed at room temperature for 15-20 minutes then washed 2 times for 5 minutes each in PBS + 0.05% tween (PBS-T) to eradicate any cryoprotectant. Each brain section was encircled with an immuno pen. After that, sections were fixed in ∼100ul of 4% paraformaldehyde (PFA; Thermo Fisher Scientific, Rockford, IL, USA) at room temperature for 15 minutes and then washed 2 times for 5 minutes each. For 1 hour at 37 °C, these sections were put in 2N HCl, and then to neutralize the HCl, they were put in a sodium borate buffer for 30 minutes. Using PBS-T, the sections were washed 3 times for 5 minutes, and then, for 1.5-2 hours at room temperature, blocked and permeabilized in a 10% BSA solution with Triton-X (Thermo Fisher Scientific, Rockford, IL, USA). Subsequently, the sections were washed 3 times for 5 minutes and incubated overnight at 4 °C with ∼100ul primary antibodies diluted in 1% BSA+Triton-X (Thermo Fisher Scientific, Rockford, IL, USA). The following morning, the sections were washed 3 times for 8 minutes each in PBS. After adding ∼100ul of secondary antibodies diluted in 1% BSA+Triton-X (Thermo Fisher Scientific, Rockford, IL, USA), the sections were incubated for 2.5 hours at room temperature, after which they were washed in PBS- T 3 times for 10 minutes each. In order to image the slides using a Leica microscope, a drop of ProLong Gold Antifade reagent with Dapi was added to each section (Thermo Fisher Scientific, Rockford, IL, USA). Images of these slides were compiled and analyzed in ImageJ.

Using immunocytochemistry protocol^13^, additional experiments were conducted. After removing plates from incubation and washing once with PBS, 200ul of 4% PFA was added to each section for 10 minutes. Next, cells were washed gently with PBS and then blocked for 2 hours at room temperature in 200ul of 10% BSA solution. Subsequently, the BSA was aspirated off of the slides and 100ul of primary antibody (mixed with 1% BSA) were added. Overnight, the cells were incubated at 4 °C. The following morning, the cells were washed 3 times for 5 minutes each in 1% BSA, after which 200ul of secondary antibody were added to each section. Then for 2-3 hours, the plate was incubated at room temperature. After the incubation, sections were washed 3 times for 5 minutes each in PBS. A drop of ProLong Gold Antifade reagent with Dapi was added to each section as done previously, which allowed for the slides to be imaged using a Leica microscope. These images were compiled and analyzed in ImageJ.

### Cell Viability Assays

Using an MTT assay and a previously established protocol, viability assays were conducted^14^. Cells were briefly plated at 3000 per well in a 96-well plate with 6-8 replicates per condition. Cells were allowed 24 hours for attachment, after which they were treated with varying doses of TMZ from 0, 50, 100, 250, 500, 1000 uM of TMZ using our lab’s standard dose response protocol^13^. Following 48 hours of treatment, media was removed and MTT solution was added to the cells. This MTT solution was made by first diluting MTT stock reagent at 5mg/ml in dPBS. Next, this MTT stock was diluted in fresh media at a stock:media ratio of 1:10. From the formulated mixture, 110ul was added to each well, and cells were incubated at 37 °C for 3-5 hours. The media and MTT stock solution was carefully removed after incubation without pipetting down or aspirating, to avoid the possible disturbance of any crystals that had formed. Cells were resuspended in 100 ul DMSO until the wells turned purple in color, an indication that the crystals had dissolved. The plate was left at room temperature for 10 minutes. It was then read on the plate reader at an absorbance of 570 nm; data was analyzed to find percent viability in each well.

### Cellular Transfection

In order to generate lentiviral particles, low passage X293 cells (ATCC, Manassas, VA, USA) were plated at 70% confluency based on a previously cited protocol^11^. After 6 hours, these cells were then transfected using a mix of HP DNA Transfection Reagent (Sigma Aldrich, St Louis, MO, USA) diluted in OptiMEM media (Gibco, Waltham, MA, USA) as well as second generation packaging and shRNA plasmids (Dharamazon), according to the manufacturer’s instructions. After maintaining the transfected X293 cells in culture for 48-72 hours, the supernatant containing the virus was harvested. This supernatant was sterilized with a 45-micron filter and ultracentrifuged at 13,3897 RCF for 2 hours. The resulting viral pellet was resuspended in PBS and aliquoted for future use.

### Viral Transduction

Using a previously optimized protocol, we resuspended cells in a small volume (∼50 ul) of media and added ∼10-20 MOI lentivirus amounts per sample^15, 16^. Next, this virus-media mixture was spun for 2 hours at 37 °C at 850 RCF, after which these cells were plated and maintained in culture with regular media changes for 48-72 hours. In order to assess the efficiency of the resulting conditions, western blots were used.

### Western Blotting

In accordance with the protocol, cells were treated, detached using trypsin, washed with PBS, and resuspended in mammalian protein extraction reagent (M-PER; Thermo Scientific, Rockford, IL, USA)^17^. M-PER was supplemented with protease and phosphatase inhibitors (PPI; Thermo Scientific, Rockford, IL, USA). After aggressively vortexing these cells for 3x 1-minute increments, with 10 minutes of rest on ice between each vortexing, the resulting lysates were spun at 13,000 RPM for 10 minutes in a 4 °C. The supernatant was then collected, and the protein concentration for each western blot was specified via BCA assay (Thermo Scientific, Rockford, IL, USA). Each sample was composed of equal amounts of protein in addition to varying amounts of sodium dodecyl sulfate buffer (SDS sample buffer; Alfa Aesar, Ward Hill, MA, USA) supplemented with beta-mercapto-ethanol and water, which allowed for each sample to contain the same total volume. After each sample was mixed, they were boiled at 95 °C for 10 minutes.

After running through 8% SDS-polyacrylamide (SDS-PAGE; made in house) by gel electrophoresis (BioRad, Hercules, CA, USA), the proteins were transferred, using a transfer machine, onto 0.45 polyvinylidene difluoride (PVDF) membranes (Millipore, Darmstadt, Germany). Following the transfer, these membranes were washed 3 times for 10 minutes each in PBS and then blocked with Tris-buffered saline (TBS) consisting of 5% powdered milk and 0.05% Tween20 (Sigma Aldrich, St. Louis, MO, USA). This blocking lasted 2 hours, after which the membranes were cut according to the proteins of interest. Next, the membranes were placed into primary antibody solutions that contained the appropriate ratio of antibody to 5% BSA solution supplemented with sodium azide, and incubated overnight on a shaker at 4°C. The following morning, these membranes were washed and then incubated for 2 hours in secondary antibody diluted 1:4000 in 5% milk. Subsequently, the membranes were washed 3 times for 10 minutes each and coated with enhanced chemiluminescence (ECL; Clarity ECL, BioRad). Using X-ray film, images were developed.

### Immunoprecipitation

A mixture of 3-5 µl of rabbit IgG antibodies and 100 µg of protein sample against ubiquitin, brought to a final volume of 250 µl with MPER buffer was incubated to produce beads according to the immunprecipation protocol we follow^18^. Samples were stored in eppendorf tubes and placed in a rotary shaker in a cold room overnight. The following day, 30 ul of the beads were administered to the IP reaction. Samples continued to be rotated for 2 hours at room temperature. Tubes were centrifuged for one minute at 1000 RPM, and supernatant was discarded. Beads were washed 3 times with 1 mL of 0.2% TBST wash buffer and centrifuged at 1000 RPM for one minute. This procedure was carried out 3 times, with the supernatant being discarded preceding each repetition. After the third and final wash, elution buffer was warmed in a 47°C water bath, 50 ul of elution buffer was added to samples, samples were resuspended, and incubated at 55°C for 10 min to elute protein from beads. After being incubated at 55 C for 10 minutes, recovered proteins were centrifuged at 3,200 RCF for 5 minutes to dissociate the beads from the supernatant. The supernatant was collected, and the aforementioned elution process was repeated. Tubes were centrifuged again at 3,200 RCF for 5 minutes. The supernatant was added to the previously new labeled tube to reach a volume of 100 ul of supernatant. 10 ul of 1 M NaOH neutralization buffer was added to each sample. Then, 28 ul of 4X SDS was added to each sample. The samples were then boiled at 95°C for 10 minutes. Western blots were then run for the analysis of precipitated proteins.

### Flow Cytometry

Cells were collected in a 96-well V bottom plate and spun down to a pellet. Cells were then washed with 100 µl of PBS. Primary antibodies (in FACS buffer, 50 ul per well) were added for one hour at room temperature in darkness. Cells were then spun down into a pellet and washed once again with 100 µl PBS. The cells were then resuspended in 80 µl of fluorescence activated cell sorting analysis (FACS) buffer. Cells were spun down again, fix perm buffer was prepared (1 part fix/perm, 3 parts buffer), and 100 ul of fix perm was added to each well. Samples were incubated for 20 minutes at room temperature in darkness. 1:10 diluted perm buffer was prepared in ddH20 and 100 ul of perm buffer was added on top of fix/perm. Samples were incubated for 10 minutes in darkness at room temperature. Samples were spun down for 5 minutes at 1,500 RPM and washed with 1:10 perm buffer once. Primary antibody was added in perm buffer (50 ul per well) and incubated overnight at 4 degrees. Samples were then washed 3 times with FACS buffer. Secondary antibody was added in FACS buffer for 1 hour at room temperature in darkness. Samples were washed and resuspended in 100 ul FACS buffer. Samples were then analyzed.

### Metabolomics isolation and liquid chromatography-mass spectrometry profiling

To measure the relative abundances of specific intracellular metabolites, extracts were prepared and analyzed by LC-MS/MS, as described previously^19, 20^. For targeted steady-state samples, metabolites were washed with 4-mL PBS and extracted on dry ice with 4-mL 80% methanol (−80°C), as described previously (Weinberg et al., 2019; Yuan et al., 2019). Insoluble material emerged as a pellet by centrifugation at 3,000 RCF for 5 min, followed by two consecutive extractions of the insoluble pellet with 0.5 mL 80% methanol, with centrifugation at 20,000 RCF for 5 min. The 5 mL metabolite extract from the pooled triplicate supernatants was dried down under nitrogen gas using the N-EVAP (Organomation, Inc, Associates). 50% acetonitrile was added to the samples, followed by vortexing for 30 seconds. Sample solutions were then centrifuged 20,000 RCF for 30 min at 4°C. Supernatant was collected for LC-MS analysis.

### Hydrophilic metabolite profiling

For complete metabolomic profiling, samples were analyzed by high-performance liquid chromatography, high-resolution mass spectrometry and tandem mass spectrometry (HPLC-MS/MS) that our lab previously has performed^19^. The system consisted of a Thermo Q Exactive^TM^ with an electrospray source, and an Ultimate 3000 (Thermo) series HPLC which consisted of a binary pump, degasser, and auto-sampler outfitted with an Xbridge Amide column (Waters; dimensions of 4.6 mm 100 mm and a 3.5 mm particle size). The mobile phase A contained 95% (vol/vol) water, 5% (vol/vol) ACN, 20 mM ammonium hydroxide, 20 mM ammonium acetate, pH 9.0; B was 100% ACN. The gradient was as follows: 0 min, 15% A; 2.5 min, 30% A; 7 min, 43% A; 16 min, 62% A; 16.1–18 min, 75% A; 18–25 min, 15% A with a flow rate of 400 ll/min. The capillary of the ESI source was set to 275C, with sheath gas at 45 arbitrary units, auxiliary gas at five arbitrary units, and the spray voltage at 4.0 kV. In positive/negative polarity switching mode, an m/z scan range from 70 to 850 was chosen, and MS1 data were collected at a resolution of 70,000. The automatic gain control (AGC) target was set at 1 x 106, and the maximum injection time was 200 ms. The top five precursor ions were subsequently fragmented, in a data-dependent manner, using the higher energy collisional dissociation (HCD) cell set to 30% normalized collision energy in MS2 at a resolution power of 17,500. Data acquisition and analysis were performed using Xcalibur 4.1 software and Tracefinder 4.1 software, respectively (both Thermo Fisher Scientific).

### 3H-guanine and 3H uridine incorporation into DNA

Treat cells were labelled with 1 μCi of either 3H-uridine or 3H-guanine as previously described^19^. Cells were harvested, and DNA was isolated using Allprep DNA/RNA kits according to the manufacturer’s instructions and quantified using a spectrophotometer. Next, 70 μl of eluted DNA was added to scintillation vials, radioactivity was measured by liquid scintillation counting and then normalized to the total DNA concentrations. All conditions were analyzed with biological triplicates and representative of at least two independent experiments.

### Enrichment Mapping

Enrichment mapping was performed using gProfiler and Cytoscape. Genes of interest were first inputted into gProfiler and the appropriate enrichment functional pathways were selected. The outputted GEM file was then uploaded into Cytoscape using the EnrichmentMap extension. This generated a base EnrichmentMap with enriched pathways and their corresponding significance (p-value). Pathways were grouped and represented as a figure.

### Statistics

All statistics were performed using the analysis software indicated or by use of GraphPad Prism software version 9.0. T-tests and ANOVAs (one- and two-way) were used to perform analyses and all P-values reported are adjusted during multiple comparisons analysis.

## RESULTS

### Single-cell RNA Sequencing Screen Identifies Uniquely Expressed Genes During TMZ Therapy in GBM

In order to investigate GBM chemoresistance and consequent tumor recurrence, our lab designed an *in vivo* experiment that utilized single-cell RNA sequencing to profile transcriptome expression and epigenetic shifts of individual cells before, during, or after TMZ-therapy. Our sequencing screen was developed in order to identify and target specific genes and networks of genes that are active during TMZ therapy to find novel pathways underlying TMZ resistance. In our model, mice treated with our vehicle control DMSO represent ‘primary GBM’, mice treated with TMZ represent ‘recurrent GBM’, and mice that received two of the five treatments of TMZ and then euthanized represent ‘during therapy GBM’ (Figure 1A). After sample preparation, 200,000 GBM43 cells were passed through droplet-based sequencing to then be sent to scRNA-seq (Figure S1a). Using Seurat Analysis and Principal Component Analysis, we identified 15 distinct clusters from our screen (Figure S1b). For better cluster visualization, further dimension reduction was performed through Uniform Manifold Approximation and Projection (UMAP) (Figure 1B). Next, we highlighted the distribution of cells based on treatment condition within the UMAP projections of our scRNAseq data. Interestingly, cells sequenced during TMZ therapy were uniquely isolated in their transcriptome profile, compared to cells sequenced in recurrent or primary GBM conditions (Figure 1C). We next wanted to determine whether the treatment specific populations correspond directly to certain phases of cell cycle. We found no significant enrichment of any cell cycle phase specific to a certain treatment condition (Figure 1D).

**Figure 1:**
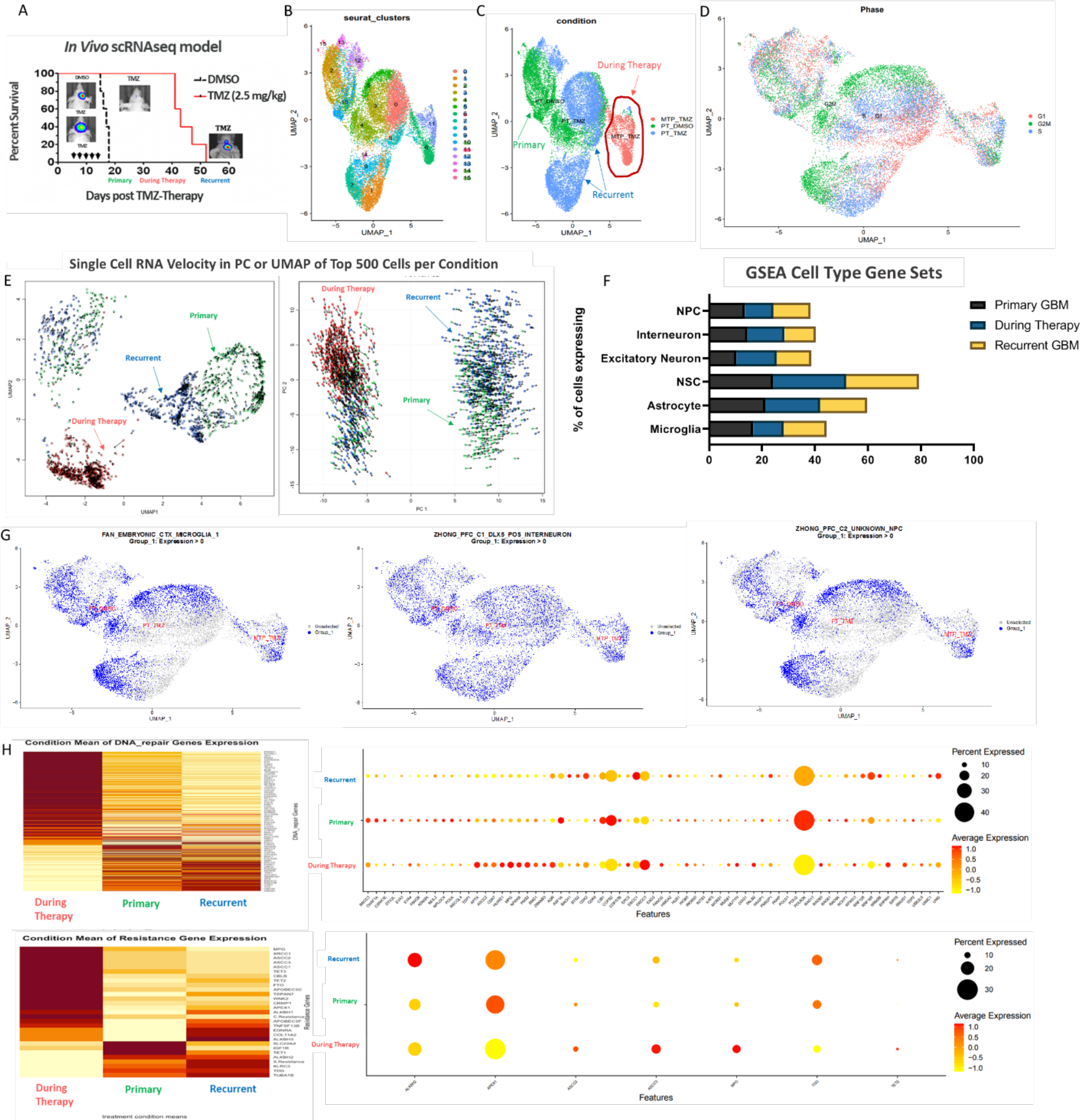
Single-cell RNA sequencing screen identifies uniquely expressed genes during TMZ therapy in GBM. **(A)** Schematic diagram of the experimental procedure. Mice given 5 treatments of DMSO (2.5 mg/kg IP) are representative of the ‘primary tumor’ model, whereas mice given 5 treatments of TMZ (2.5 mg/kg) during treatment are representative of the ‘recurrent tumor’ model and both groups are taken for scRNA-seq at survival endpoint. Mice taken for scRNA-seq in the ‘during therapy’ received 2 of the 5 TMZ treatments and were then euthanized. **(B)** Seurat clusters created via UMAP dimension reduction of all scRNA-seq quality-controlled data from all samples of all three groups. **(C)** Visualization of group distribution within cluster data. **(D)** Visualization of cell cycling phase within cluster data. **(E)** RNA velocities were computed via velocyto^21^. This includes the scRNA-seq data from the top 500 quality-controlled cells of the three groups. The RNA velocity data is shown using UMAP reduction (left) and PCA reduction (right). **(F)** GSEA bar graphs of cell type gene sets expressed in our scRNAseq data. **(G)** UMAP cell distribution of cell type gene sets expressed in our scRNAseq data. **(H)** GSEA heat maps and dot plots of DNA repair associated genes (above) and resistance associated genes (below) separated by group condition expressed in our scRNAseq data.

To further investigate the unique genetic expression profile of cells, present during therapy, we used velocyto with both UMAP and PCA reduction techniques to analyze the single-cell RNA velocity of each cell^21^. RNA velocity plots use high-dimensional vectoring analysis of spliced, unspliced, and degraded mRNA to showcase active transcription and emerging genetic profiles o single cells. We then highlighted which treatment conditions the cells belonged to within our RNA velocity graphs. Similar to the clustered cell distribution illustrated when mapping transcriptomic data by treatment condition, velocity plots presented a distinct cluster of RNA velocities, including specific vector size and direction, from cells sequenced during TMZ therapy (Figure 1E). This suggests that during TMZ therapy cells are in a unique transcription state in comparison to both primary and recurrent tumor transcription. Our *in vivo* scRNA-seq pipeline was developed in order to identify and target specific genes and networks of genes that are active during TMZ therapy to find novel pathways underlying TMZ resistance.

Identifying the expression profile of upregulated genes during and post therapy, allowed us to further explore the differences between two treatment conditions’ populations (at this point you haven’t actually discovered what the expression profile is yet). We first used Gene Set Enrichment Analysis (GSEA) to determine the presence of various cell types in our data, in which we see the cell type distribution to be relatively consistent across treatment conditions (Figure 1F, Figure 1G). We next wanted to use GSEA to see how Oncogenic Markers present in our data. Here, we can see a drastic difference in both the expression of DNA Repair Gene Expression and Resistance Gene Expressions as they are distributed in our data heat map. As you can see, there is a visual distinction between genes expressed during therapy compared to primary and recurrent GBM (Figure 1H).

### Computational and Experimental Validation of Oncogenic Gene Sets in Sequencing Data

We next wanted to determine how published oncogenic gene sets appear in our scRNA-seq data to explore how we could experimentally validate our computational data. If we examine gene sets that present the highest mean expression of in our sequencing data during therapy, for example KRAS.50_UP.V1_UP, we can not only visually confirm this enhanced expression through feature plots and dot plots in which the KRAS gene set is elevated during therapy, but we can map the enriched pathways of the gene sets, using gProfiler and Cytoscape, and see that functionally, the highly expressed genes cluster into pathways underlying “Response to Chemical,” “Chemotaxis,” “Regulation of Biological Quality,” etc., suggesting the enriched pathways contribute to chemoresistance during therapy (Figure 2A). We see the same pattern when we analyze the gene set expression of ALK_DN.V1_UP which shows to be elevated in expression during therapy and functionally is involved in pathways such as “Response to External Stimulus and Stress” and “Response to Organic Substance,” indicating a strong functional response during therapy (Figure 2B). In contrast, when we evaluate the expression of oncogenic gene sets that are elevated in recurrent GBM, such as RB_P107_DN.V1_UP and MTOR_UP.V1_UP, we can see high expression in our scRNA-seq recurrent model, as well as a functional shift in enriched pathways correlating to homeostatic processes such as metabolic processes and cellular organization, indicating adaptive mechanisms to regain proliferative functions post-therapy (Figure 2C).

**Figure 2:**
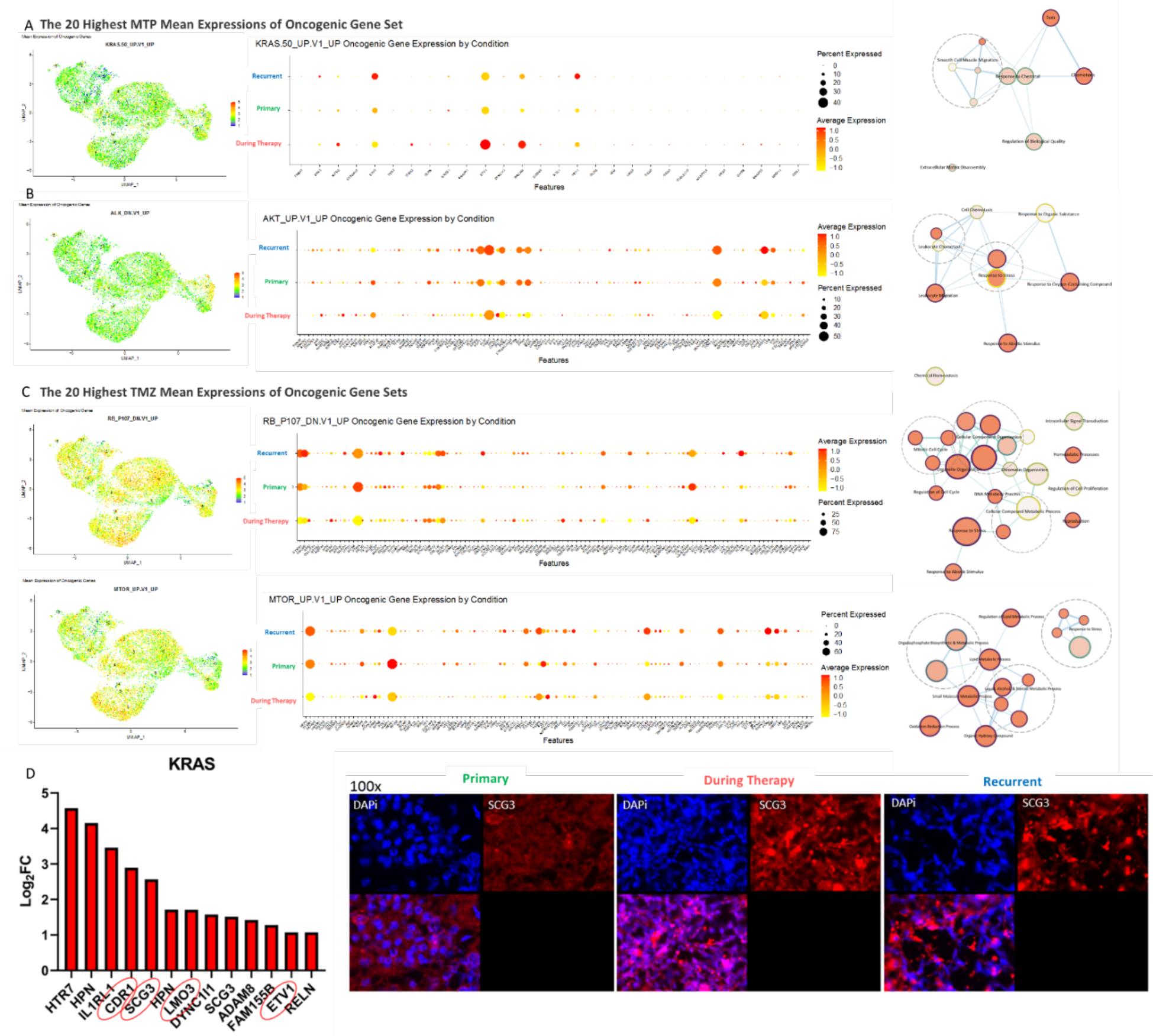
Computational and experimental validation of oncogenic gene sets in sequencing data. **(A-B)** The top two uniquely highly expressed oncogenic genes sets found in GSEA of our during therapy scRNA-seq model. KRAS.50_UP.V1_UP (above) refers to the set of oncogenes upregulated in tumor cell line overexpressing KRAS. ALK_DN.V1_UP (below) refers to the set of oncogenes upregulated in tumor cell line with KD of ALK. The distribution of the GSEA enrichment was overlayed onto the Seurat clusters for all quality-controlled samples of all three conditions (left). The GSEA enrichment was compared between all three conditions for each gene included in the gene sets (middle). Pathway analysis of oncogenic gene sets was performed using gProfiler and Cytoscape (right). **(C)** The process as described above was repeated but for the top two uniquely highly expressed oncogenic gene sets found in GSEA performed on our recurrent scRNA-seq model. RB_P107_DN.V1_UP (above) refers to the set of oncogenes upregulated in RB1 and RBL1 KO mice. MTOR_UP.V1_UP (below) refers to the set of oncogenes upregulated with everolimus (RAD001) treatment, an MTOR inhibitor. **(D)** The apparent log fold-change expression of KRAS gene set in our RNA-seq data when comparing during therapy to recurrent GBM conditions. Genes circled were chosen for later verification of GSEA data due to their high enrichment during therapy as well as their apparent protein interactions within the KRAS gene set. **(D)** Immunohistochemistry of primary, during therapy, and recurrent GBM tissue and stained for nucleus in blue (DAPi) and SCG3 in red (APC).

Lastly, we can experimentally validate the distinction between cellular populations present during therapy compared to primary and recurrent GBM. We mapped the enriched genes of KRAS.50_UP.V1_UP and ALK_DN.V1_UP onto our sequencing data in order to quantify which genes of these two oncogenic gene sets would be the most upregulated in our scRNA-seq model of during therapy. For example, in the KRAS gene set, we can see SCG3 is highly elevated, and using immunohistochemistry (IHC) in tissue of primary, during therapy, and recurrent, we can see that SCG3 is in fact elevated during therapy – experimentally validating our single cell and computational analysis (Figure 2D).

### Identified Genes Show Significance in Pathway Analysis

Computational analysis and complementary immunohistochemistry strongly suggested that the population of cells present during therapy was distinct from cells in primary and recurrent GBM. In order to investigate mechanisms underlying chemoresistance in GBM, our next goal was to validate targets of interest from our single-cell sequencing data. Using a combination of gProfiler and Cytoscape, we mapped the pathways associated with genes significantly elevated during therapy. Of several enriched pathways, ‘Metabolic Processes’ (*p<0.00001*) particularly interested us. We then used Stringr to map gene networks involved in metabolic processes found to be elevated in our sequencing data. The RNR family of genes (RRM1, RRM2, and RRM2B), essential to dNTP metabolism, exhibited a unique signature in our sequencing data (Figure 3A). During our pathway analysis, the enriched pathways in recurrent GBM as well as the depleted pathways during therapy and in recurrent GBM were also mapped (Figure S2a-b).

**Figure 3:**
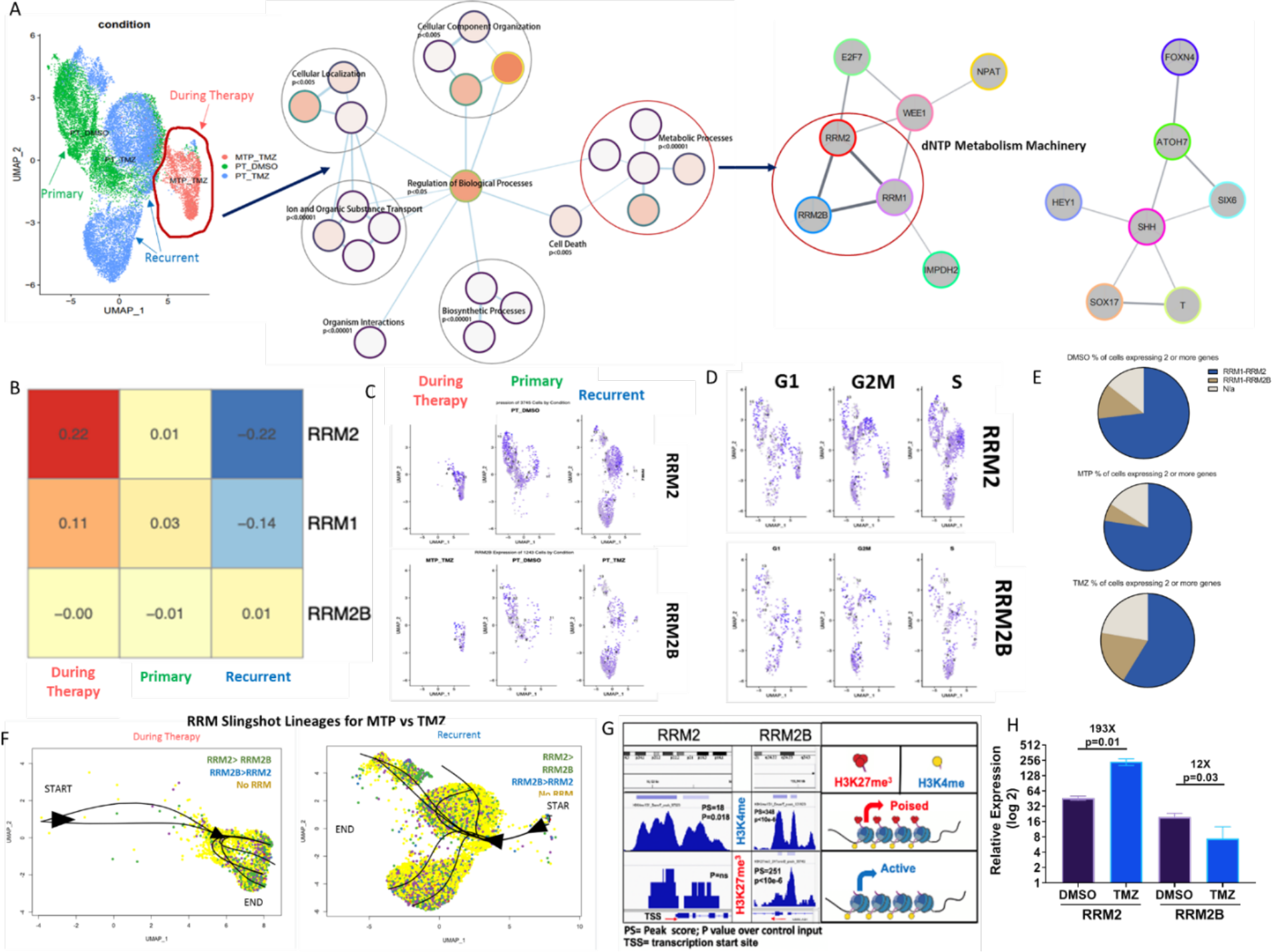
Identified genes show significance in pathway analysis. **(A)** Schematic of how RNR genes were identified for future analysis. Metabolic genes were found to be among the highest enriched processes in our during therapy scRNA-seq data, which was further delineated into RNR genes as well as other metabolic genes. RNR genes were taken as a starting point for further investigation from this analysis. **(B)** Representative heat map of the enrichment of specific RNR genes across all three conditions. RRM2 was found to have the most uniquely upregulated expression during therapy, followed by RRM1, and then RRM2B which had no significant difference. **(C)** The complete Seurat cluster data was highlighted for cells that were present during therapy (left), in primary GBM (middle), or in recurrent GBM (right). Cell distribution was mapped for high RRM2 expression (above) or RRM2B (below). **(D)** The complete Seurat cluster data was highlighted for cells that were in certain cell cycle phases, G1 (left), G2-M (middle), or S (right) and expressed either RRM2 (above) or RRM2B (below). **(E)** A pie chart showing the apparent expression of both RRM1 and RRM2 (blue) or RRM1 and RRM2B (dark brown) in cells that originate from each scRNA-seq condition. **(F)** Seurat clusters were created using only during therapy scRNA-seq data (left) or only recurrent GBM scRNA-seq data (right). The lineage progression of possible cell development from cluster to cluster was determined using the program Slingshot. The progression of the lineages is shown using the black line projections from the starting cluster. Cells were then highlighted by their expression status, with either expression of RRM2>RRM2B, RRM2B>RRM2, or no significant RNR gene expression. **(G)** Schematic of whole genome ChIP-seq analysis for H3K4me1 and H3K27me3 at the TSS of RRM2 and RRM2B. **(H)** mRNA expression of RRM2 or RRM2B in conditions of DMSO or TMZ analyzed through RNA-seq.

In our single-cell sequencing analysis, while RRM1 and RRM2B remain relatively constant in primary, during therapy, and recurrent sequencing data, RRM2 is significantly upregulated during therapy (Figure 3B). A UMAP blot of the raw distribution of cells expressing high levels of either RRM2 or RRM2B in primary, during therapy, or recurrent GBM shows RRM2 cells to be the most concentrated during therapy. These feature plots confirm the original heat map signature displayed, in which while RRM2B remains constant, RRM2 expression is elevated during therapy (Figure 3C). When we map the cells expressing high levels of RRM2 or RRM2B onto their cell cycle phases, we can see that again, while RRM2B remains constant, there are higher levels of RRM2 in the S and G2M phase, indicating enhanced proliferation and cell growth (Figure 3D). We then calculated the percentages of cells expressing high levels of RRM1 and RRM2, compared to high levels of RRM1 and RRM2B to model expression changes RRM1-RRM2 throughout TMZ therapy. Our data suggests of all cells expressing RRM1, RRM2, RRM2B, the cells expressing high levels of RRM1-RRM2, increases from 58.8% in recurrent GBM to 91.4% in GBM cells during therapy (Figure 3E, Figure S2c). Another way to visually determine the preference of RRM2 over RRM2B during therapy, was to map the Slingshot lineages of cells expressing the RNR gene family in both during therapy and recurrent GBM. Analyzing our single cell RNAseq data through Slingshot allowed us to create distinct lineages of cellular populations as they evolve throughout pseudotime, to model the developmental trajectory sequencing data^22^. Compared to recurrent GBM, it is evident that the trajectory of cells during therapy can be predicted to evolve into cells in which RRM2 expression is higher than RRM2B (Figure 3F).

Finally, using only one screen to find novel targets to pursue has its limitations, so next we wanted to validate this family of genes against other screens that our lab has performed in the past. Using our CHIPseq screen in which we sent a whole genome chip sequencing of Day 1 vs. Day 4 of TMZ therapy in order to analyze 3 different markers, our analysis reveals that RRM2B has upregulated H3K4 monomethylation, as well as upregulated H3K27 trimethylation. These two markers upregulated in combination both activate and repress transcription causing the histone to be poised and temporarily stopped. We also see that RRM2 has upregulated H3K4 monomethylation which means that there is an immediate increase in binding at the transcription start site (Figure 3H). Our lab intentionally sent our CHIPseq and a basic DMSO vs TMZ RNAseq at the same time in order to correlate the effect of transcription in real time. We show that in TMZ, RRM2 is upregulated at the transcription start site (TSS) in CHIPseq and our RNAseq confirms that RRM2 is in fact upregulated. In contrast, we show that in TMZ, RRM2B transcription was temporarily paused in CHIPseq and our RNAseq presents RRM2B to be downregulated in TMZ (Figure 3I). Taking all of this into consideration lead us to believe that the RNR family of genes, especially RRM2, is incredibly important during therapy, so we decided to pursue them further.

### Ribonucleotide Reductase Family Genes Show Effects on Patient Survival

To better understand how RNR family gene expression correlates with survivability in GBM, we examined publicly available GBM patient datasets. We first compiled the mutation profiles of RRM2 and RRM2B genes using cBioPortal, which revealed that mutation rates were below 2% across 248 patient samples (Figure 4A). Next, using a public GBMseq analysis of neoplastic cells in GBM patients, we identified cells expressing significant levels of RRM2 to be concentrated in the tumor core, whereas cells expressing RRM2B are distributed among both the tumor core and periphery, as measured through log2counts per million (Figure 4B). Since RRM2 is upregulated during therapy in our scRNA-seq, we expect it to be correlated with chemo-resistance and decreased patient survival. Using Kaplan Meier Survival Estimates, we see that high RRM2 expression is correlated with lower survival (*p=0.0012*) while high RRM2B expression does not have a significant effect on survival (Figure 4C). Next, through the GlioVis portal, we analyzed both The Cancer Genome Atlas (TCGA) and the Chinese Glioma Genome Atlas (CGGA) data of RRM2 and RRM2B. We found that RRM2 mRNA expression is significantly upregulated in GBM samples compared to non-tumor samples, whereas RRM2B mRNA expression is relatively equal between the two sample populations (*p<0.001*). Additionally, RRM2 mRNA expression was found to significantly increase with tumor grade from grade II to IV (*p<0.001*), while RRM2B mRNA expression does not significantly change based on tumor grade. We also see that mRNA expression for both RRM2 and RRM2B is significantly correlated with classical and mesenchymal GBM tumor subtypes (*p<0.001*) (Figure 4D, Figure S3a-b). Using GlioVis databases to generate correlation plots between RRM1, RRM2, and RRM2B, we can see that RRM1-RRM2 is positively correlated in GBM (*p=0.00*), while RRM1-RRM2B is negatively correlated (*p=0.00*) (Figure 4E, Figure S3c). To explore the relationship between the mRNA expression of our genes and tumor recurrence status, we used The Human Protein Atlas to examine RNR protein expression within recurrent glioma tissue sample. RRM2 consistently shows a low (<25%) expression stain, while RRM2B shows a high (>75%) expression stain in post-therapy tumor tissue which aligns with our scRNA-seq data (Figure 4F). Finally, we performed a western blot that included samples from cancer vs. non-cancer cell lines (Fibroblast, Astrocyte, H1B, Neural Stem Cell, GBM43, GBM6, GBM39, SNB-19, and U251) and stained for RRM1, RRM2, and RRM2B in order to compare expression differences between cell lines and confer the baseline clinical expressions of RNR proteins. This blot revealed that RNR gene expression is relatively higher in PDX GBM cell lines compared to astrocytes, fibroblasts and neural stem cells; however, between PDX lines there are differences in expression (Figure 4G). To account for GBM subtype differences and subtype-specific mechanisms, we performed our experiments in multiple PDX cell lines.

**Figure 4:**
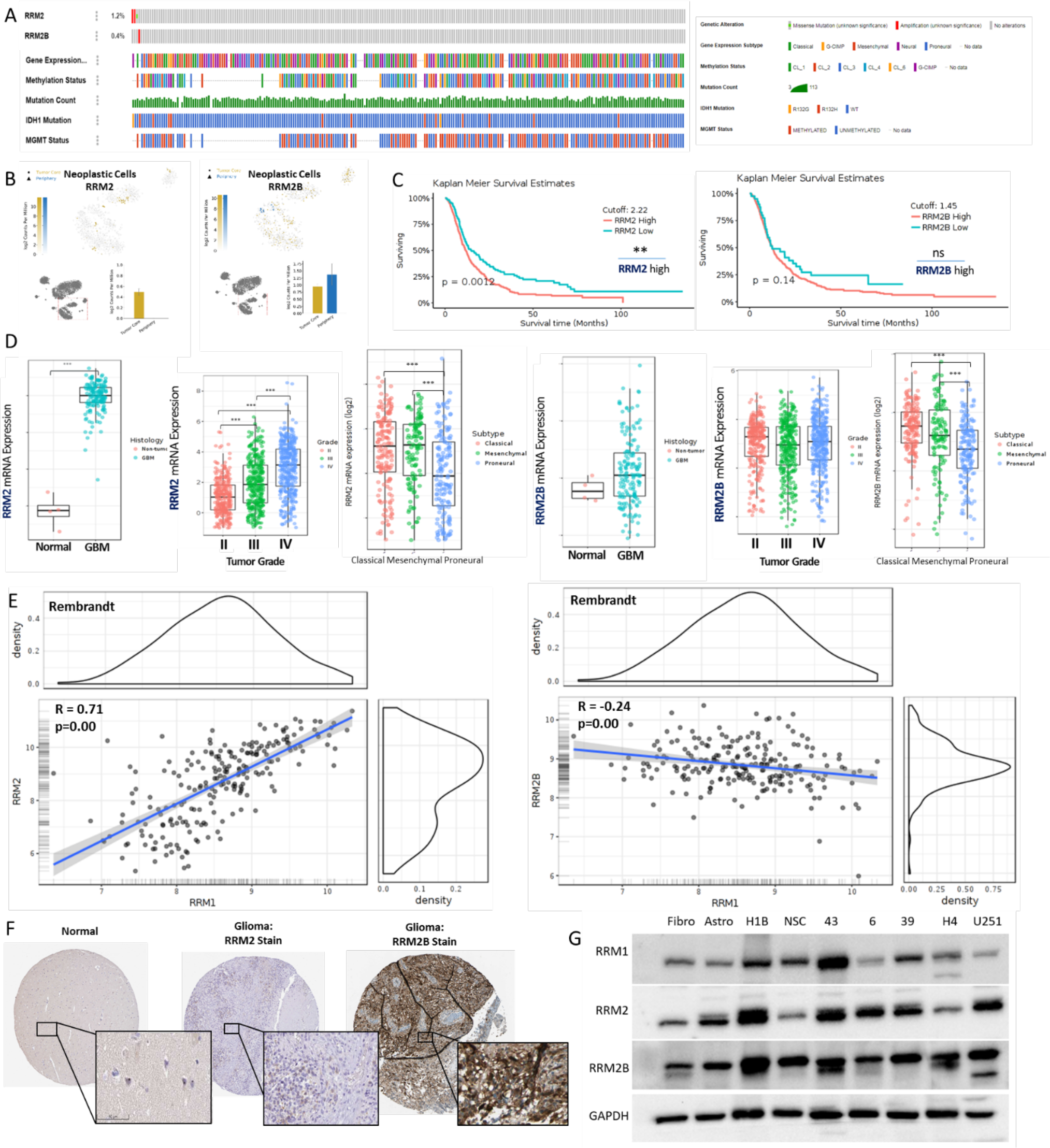
Ribonucleotide Reductase family genes show effects on patient survival. (A) Mutation rates of RRM2 and RRM2B genes were below 2% across 248 patient samples as identified through cBioPortal. Corresponding key is located to right of mutation plot. (B) Expression of RRM2 (left) or RRM2B (right) in tumor core or tumor periphery. (C) Kaplan Meier Survival Estimates identify high RRM2 expression (left) to be correlated to lower patient survival, while RRM2B expression (right) does not have significant correlation with patient survival. (D) Data from GlioVis acquired through TCGA and CGGA databases found RRM2 mRNA expression (left) to be significantly upregulated in GBM tumor tissue and in higher glioma tumor grade, while RRM2B mRNA expression (right) is not significantly expressed in tumor tissue or correlated with higher grade gliomas. Both RRM2 (left) and RRM2B (right) mRNA expression is higher in classical and mesenchymal tumor subtypes. (E) Rembrandt Correlation plots between RRM1-RRM2 (left) and RRM1-RRM2B (right) generated on GlioVis. (F) The Human Protein Atlas identifies RRM2 protein expression to be <25% in recurrent tissue, while RRM2B expression is >75%. (G) Baseline expression profiles of genes of interest in all cell lines used in this study. *p < 0.05; **p < 0.01; ***p < 0.001; ****p < 0.0001; ns, not significant.

### RNR Gene Expression Altered During TMZ Therapy

After exploring the clinical significance of RNR family genes, we conducted a literature review to better understand the research that has been done on the ribonucleotide enzyme in cancer. Biologically, beta subunits RRM2 and RRM2B share 81% sequence homology. Using NCBI Protein BLAST, we show the sequence alignment of RRM2 and RRM2B (Figure 5A). Functionally, it is thought that subunit RRM2 is actively expressed in proliferating cells, whereas RRM2B is only required for non-proliferating cells^23^. Foskolou et al. (2017) hypothesized that in hypoxic conditions, the ribonucleotide reductase complex requires subunit switching to maintain the integrity of DNA replication of colon carcinoma cells in hypoxic conditions^24^. While GBM patient datasets and survival statistics demonstrated the clinical significance of differential RNR family gene expression, the functional roles of RRM2 and RRM2B and subunit binding preferences have yet to be addressed in the context of GBM. We first established how RNR proteins are expressed in GBM cell lines during TMZ therapy *in vitro*. After multi-exposure treatments of TMZ or equimolar vehicle control DMSO over the course of 72 hours, GBM PDX cell lines maintain constant expression of RRM1 and RRM2B while RRM2 increases with TMZ exposure. Western blots performed in GBM6 and GBM43 show no change in RRM1 and RRM2B protein levels between DMSO and TMZ conditions or upon repeat exposure to control or TMZ-based therapy. In contrast, RRM2 protein concentrations increase from 1x to 3x exposure to chemotherapy but remain unchanged across corresponding DMSO multi-exposure (Figure 5B, Figure S4a). Next, to validate the interaction between the alpha and beta subunits of the RNR complex. We subjected protein from GBM cell lysates to immunoprecipitation with an anti- RRM1 antibody and performed subsequent western blotting which confirmed RRM1-RRM2 and RRM2-RRM2B interactions across treatment conditions. DMSO conditions exhibit an equilibrium between RRM1-RRM2 and RRM1-RRM2b interactions between 48h and 96h. Our IP-WB data shows preferential RRM1-RRM2 interactions in the early post-treatment time point at 48h compared to the later post-treatment time point at 96h which supports our scRNA-seq finding of enriched RRM2 transcripts during therapy compared to recurrent tumors (Figure 5C). We used fluorescence-activated cell sorting to confirm our IP-WB findings in multiple PDX cell lines. Our data shows a higher percentage of GBM cells expressing RRM2 during TMZ therapy compared to control treatment. RRM2B expression during TMZ therapy also increases slightly in GBM6 but remains constant compared to DMSO (*p<0.001*) (Figure 5D, Figure S4b). FACS analysis comparing RRM2 expression across two time-points showed high levels of RRM2 immediately post-TMZ at 24h (*p<0.001*), but diminished RRM2 expression after 72h thus substantiating our single cell screen results (Figure 5E, Figure S4c).

**Figure 5:**
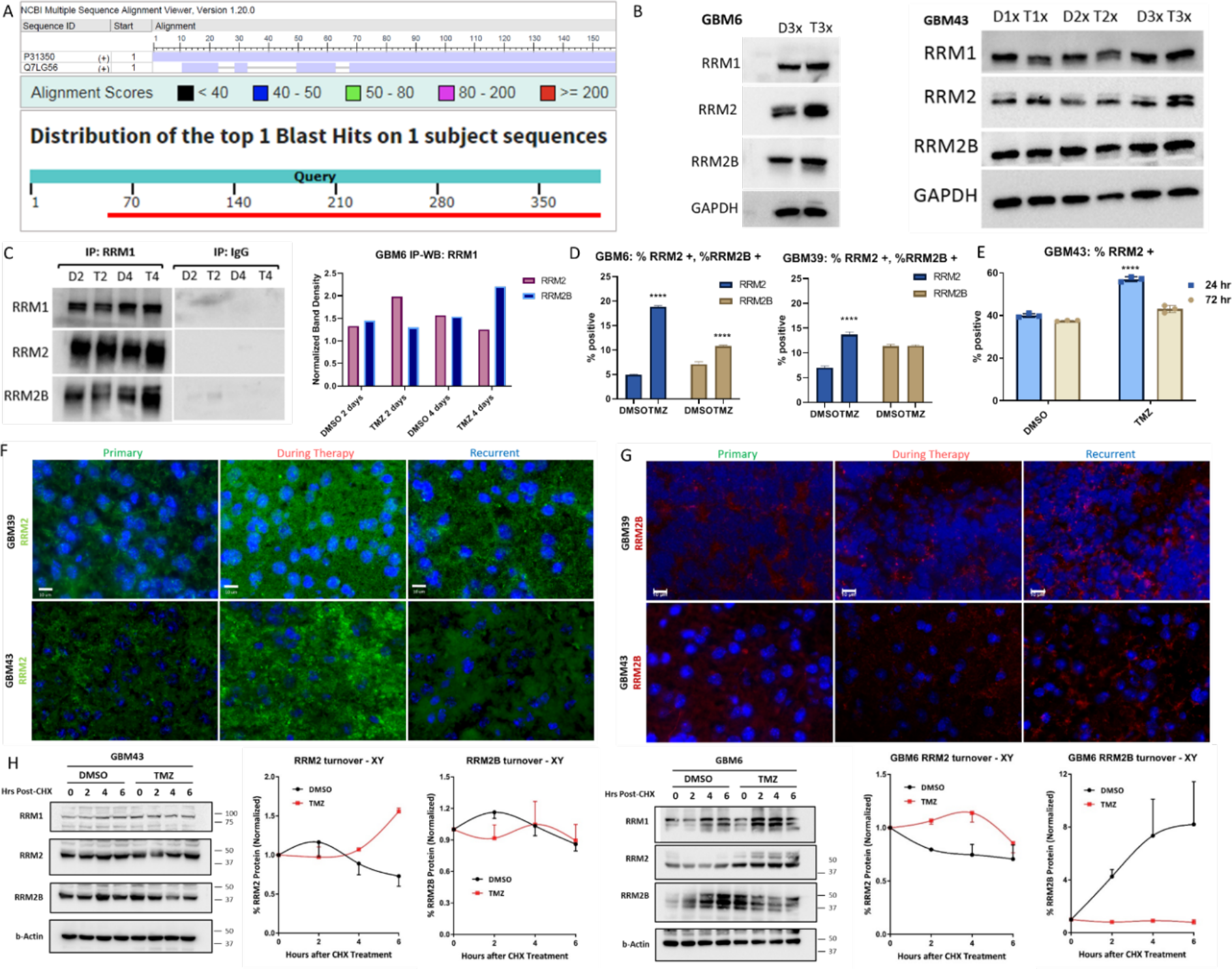
RNR gene expression altered during TMZ therapy. **(A)** NCBI Protein BLAST to show sequence alignment of RRM2 and RRM2B. **(B)** Western blot analysis of genes of interest when exposed to TMZ (50uM) or DMSO (uM) for a total of 1, 2, or 3 times (DMSO 1x : D1x). Validated in GBM6 and GBM43. **(C)** Immunoprecipitation and consequent western blot analysis of RRM2 and RRM2B binding to RRM1 in DMSO or TMZ (50uM) for 2 or 4 days (DMSO 2 days: D2). Using ImageJ bands were normalized to GAPDH and bar graphs of IP interaction were created. **(D)** Representative FACS bar graphs for intracellular RRM2 or RRM2B staining in DMSO or TMZ (50uM. Validated in GBM6 and GBM29. **(E)** Representative FACS bar graphs for intracellular staining of RRM2 in DMSO or TMZ (50uM) for 24 or 72 hours. Validated in GBM43. **(F-G)** Immunohistochemistry of primary, during therapy, and recurrent GBM tissue stained for RRM2 (FITC) or RRM2B (APC). Validated in GBM39 and GBM43. **(H)** Chase Assay and western blot analysis of cells treated with Cylcoheximide (CHX), and DMSO or TMZ (50uM), and stained for genes of interest. Corresponding graphs made in Prism 9.0 for chase assay genes of interest.*p < 0.05; **p < 0.01; ***p < 0.001; ****p < 0.0001; ns, not significant.

Immunohistochemistry on tissue from our *in vivo* scRNA-seq model of primary, during therapy, and recurrent GBM illustrates a stark difference in RRM2 expression to be significantly stronger in GBM during therapy compared to primary or recurrent GBM tissue. In parallel, RRM2B consistently exhibits low fluorescence across the same sample conditions with minimal expression across conditions and cell lines (Figure 5F). In order to understand the rate of RNR protein degradation and turnover during TMZ therapy, we determined the half-life of each RNR protein. We performed a Cycloheximide (CHX) Chase Assay that showed RRM2 protein turnover is significantly higher in cells up to 6 hours post-CHX treatment, compared to the protein turnover percentages of RRM1, RRM2B, and control, indicating increased availability of RRM2 during TMZ therapy (Figure 5G, Figure S4d). The CHX assay shows an upward trend in RRM2 turnover until endpoint in GBM43 TMZ vs. DMSO conditions and peak RRM2 turnover at 4h in GBM6 TMZ vs. DMSO conditions. RRM2B turnover data shows a dynamic equilibrium between DMSO and TMZ conditions in GBM43, and an upward trend in the GBM6 DMSO condition with no change over time in the GBM6 TMZ condition.

### RNR genes show effects on GBM viability during TMZ treatment

For our genes of interest—RRM1, RRM2, and RRM2B—we performed several *in vitro* experiments to examine the effect of gene knockdowns on GBM cell viability when treated with TMZ. RRM1, RRM2, and RRM2B knockdown cells lines were created using shRNA plasmid transfection and subsequent transduction (Figure S5a). Knockdown efficiency validated through western blot and supplemental densitometry which found sufficient knockdowns were consistently produced. After generating the knockdowns in PDX cell lines (GBM43, GBM6, and U251), MTT cell viability assays were performed on each cell line to investigate cytotoxicity upon increasing doses of TMZ. RRM2-knockdown cells demonstrated increased TMZ- susceptibility, whereas RRM1- and RRM2B-knockdown cells demonstrated significant resistance to TMZ (*p<0.001*) (Figure 6A, Figure S5b). We then used immunocytochemistry to investigate the DNA damage response (DDR) of our established cell lines, in which we found elevated yH2AX fluorescence in RRM2-knockdowns after TMZ treatment, signifying reduced DNA repair capacity compared to the control (*p<0.001*). Conversely, RRM2B-knockdowns show consistently reduced yH2AX fluorescence after TMZ treatment, indicating their increased efficiency to engage with DDR machinery and recover from therapy quicker as compared to control (*p<0.001*) (Figure 6B). We then validated this finding using FACS in which RRM2-knockdowns exhibited increased yH2AX staining up to three days after TMZ treatment, while RRM2B-knockdowns produced decreased yH2AX levels after TMZ treatment (*p<0.0001*) (Figure 6C). To determine whether the DDR of our RRM1, RRM2, and RRM2B-knockdowns was specific to TMZ, additional MTT assays were performed using either Ibrutinib or CCNU, two DNA-damage inducing drugs^25, 26^. RNR family knockdowns do not exhibit the same drug-induced phenotypes when treated with TMZ, indicating TMZ-specific phenotypes in RNR knockdown cell lines (Figure S5c-d).

**Figure 6:**
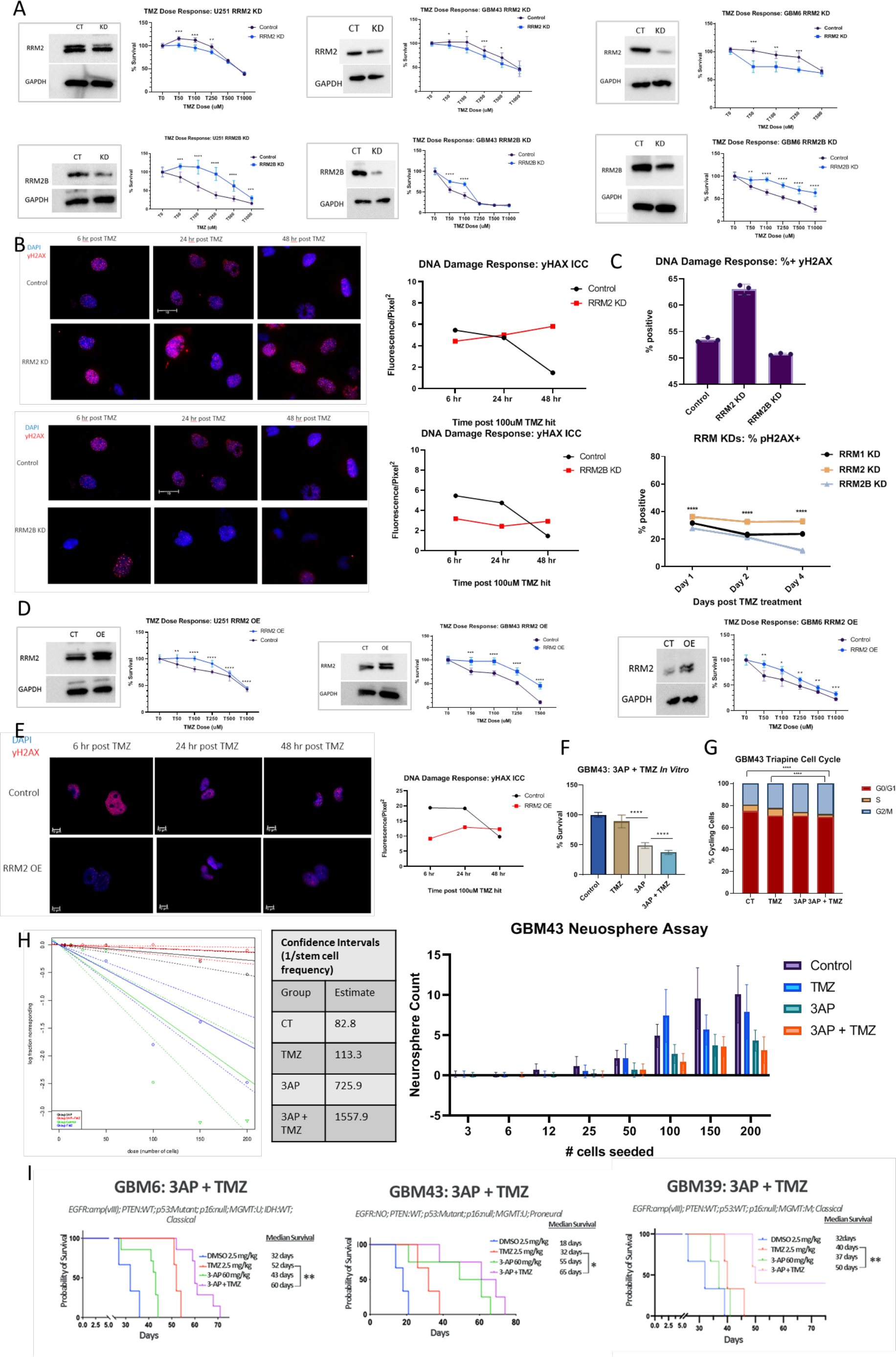
RNR genes show effects on GBM viability during TMZ treatment. **(A)** Western blot analysis of RRM2 and RRM2B shRNA KD efficiency. Cell viability assay of RRM2 KD and RRM2B KD in 48-hour TMZ dose response (0uM, 50uM, 100uM, 250uM, 500uM, 1000uM). Validated in U251, GBM43, and GBM6. Graphed in Prism GraphPad 9.0, using to compare row- means to determine significance or using log-rank tests to determine significance. **(B)** Immunocytochemistry of Control vs. RRM2 KD and Control vs. RRM2B KD cells after 100 uM TMZ treatment. Nucleus stained in blue (DAPi). yH2AX stained in red (APC). ImageJ Fiji used to quantify yH2AX fluorescence and plotted in Prism 9.0 graph. **(C)** Representative FACS bar graphs of Control, RRM2 KD, and RRM2B KD cells after 100uM TMZ treatment and intracellularly stained for yH2AX. Representative FACS line graphs of Control, RRM2 KD, and RRM2B KD cells after 100uM TMZ treatment and 1, 2, or 4 days of recovery and then intracellularly stained for yH2AX. **(D)** Western blot analysis of RRM2 OE efficiency. Cell viability assay of RRM2 OE in 48-hour TMZ dose response (0uM, 50uM, 100uM, 250uM, 500uM, 1000uM). Validated in U251, GBM43, and GBM6. Graphed in Prism GraphPad 9.0 using to compare row-means to determine significance or using log-rank tests to determine significance. **(D)** Immunocytochemistry of Control vs. RRM2 OE after 100 uM TMZ treatment. Nucleus stained in blue (DAPi). yH2AX stained in red (APC). ImageJ Fiji used to quantify yH2AX fluorescence and plotted in Prism GraphPad 9.0. **(F)** Cell viability assay of GBM43 cells treated with DMSO (50uM), TMZ (50uM), 3-AP Triapine (2uM), or TMZ + 3AP Triapine, plotted in Prism GraphPad 9.0 using ANOVA to compare row-means to determine significance or using log-rank tests to determine significance. **(G)** Representative cell cycle FACS of GBM43 cells treated with DMSO (50uM), TMZ (50uM), 3-AP Triapine (2uM), or TMZ + 3AP Triapine, stained with propidium iodide (PI), graphed in Prism GraphPad 9.0 using ANOVA to compare row-means to determine significance or using log-rank tests to determine significance. **(H)** Extreme Limiting Dilution Analysis (ELDA) plot of neurospheres form in GBM43 cells treated with DMSO (50uM), TMZ (50uM), 3-AP Triapine (2uM), or TMZ + 3AP Triapine. Corresponding table of confidence intervals generated through ELDA. Corresponding bar graph of neurosphere assay analysis plotted in Prism GraphPad 9.0. **(I)** In vivo survival analysis in mice engrafted with GBM cells and treated with DMSO (2.5 mg/kg IP), TMZ (2.5 mg/kg), 3-AP Triapine (40-60 mg/kg IP), or TMZ + 3-AP Triapine. Graphed in Prism GraphPad 9.0, using ANOVA to compare row-means to determine significance or using log-rank tests to determine survival significance. *p < 0.05; **p < 0.01; ***p < 0.001; ****p < 0.0001; ns, not significant.

Since our RRM2-knockdown experiments demonstrated the importance of RRM2 in cellular sensitivity to TMZ, we next wanted to examine the effects that overexpressing RRM2 might have on GBM cell viability. After generating RRM2-overexpression PDX cell lines (GBM43, GBM6, U251) using shRNA plasmid transfection and subsequent transduction, immunoblotting was performed to validate RRM2 overexpression in each cell line. RRM2-overexpression cell lines were calculated to each produce sufficient overexpression. MTT assays were then performed on each established cell line in which we found that in response to TMZ, RRM2-overexpression cells were resistant to TMZ. (Figure 6D). The phenotypic difference between RRM2-knockdown and RRM2-overexpression cells exhibited through TMZ dose response viability assays indicates that RRM2 is not only essential to how cells respond to TMZ but can be considered a driver of chemoresistance in GBM. To further validate this conclusion, we found RRM2-overexpression cells to show dramatically decreased yH2AX staining in response to TMZ treatment, indicating enhanced DNA repair efficiency in gliomagenesis (Figure 6E).

We then performed several *in vitro* and in *vivo* experiments to determine whether an RRM2- inhibited genotype would have the same phenotypic effect on mouse survival. A selective RRM2 inhibitor, 3-AP Triapine, which is currently FDA approved and in clinical trials as an anti-cancer agent for different human malignancies, has been demonstrated to cross the blood-brain barrier (BBB)^27^. A cell viability assay shows that 3AP + TMZ treatment, causes significantly more GBM cell death than TMZ alone (*p<0.0001)* (Figure 6F). Assessing altered cell cycle phases of GBM cells during Triapine treatment shows that compared to TMZ treated cells, 3AP + TMZ shows drastically reduced S-phase in cells, indicating Triapine’s efficiency in arresting the cell cycle and decreasing cell proliferation (*p<0.001*) (Figure 6G, Figure S5e). A neurosphere assay, to quantify stemness of GBM cells, was then performed on cells under various treatment conditions. Not only did 3AP + TMZ treated GBM cells form significantly less neurospheres compared to both TMZ and Control, but the size of said neurospheres were also significant reduced (*p<0.05*) (Figure 6H, Figure 6I, Figure S5f).

Lastly, in our *in vivo* experiments, 3-AP alone, similar to RRM2-knockdowns without treatment, does not show significant improvements on mouse survival. However, 3-AP (40mg/kg) in combination with TMZ (2.5mg/kg) significantly improves survival in mice one week post intracranial injection of GBM6, GBM39, and GBM43 cell lines (*p<0.001*). On average mice treated with 3-AP + TMZ survived significantly longer than those treated with TMZ alone (*p<0.01*). Taken together, *in vitro* RRM2-knockdowns and *in vivo* RRM2 inhibition through 3-AP Triapine, significantly sensitizes GBM cells to TMZ, indicating promising clinical opportunities in targeting RRM2 (Figure 6J).

### RRM2-mediated production of dCTP and dGTP in necessary for adaptation to TMZ therapy

To elucidate the mechanism of RRM2-mediated chemoresistance, we performed a targeted metabolomics analysis to quantify dNTP signatures of our knockdown cell lines during TMZ therapy. In response to TMZ, dCTP and dGTP production increased 100-fold and 80-fold respectfully in control cells (*p<0.01*). Interestingly, RRM2-knockdowns did not exhibit the same dNTP induction in response to TMZ, producing significantly less dCTP and dGTP pools (*p<0.001*) (Figure 7A, Figure S6a). We then preformed bulk metabolomics which allowed us to track metabolite production in control versus RRM2 KD cells throughout RNR-dependent the de novo synthesis of dNTPs. We see multiple steps in which RRM2 KD cells exhibit significantly altered metabolite production compared to controls, confirming our hypothesis that RRM2 is required for proper dNTP pool production and homeostasis (Figure 7B).

**Figure 7:**
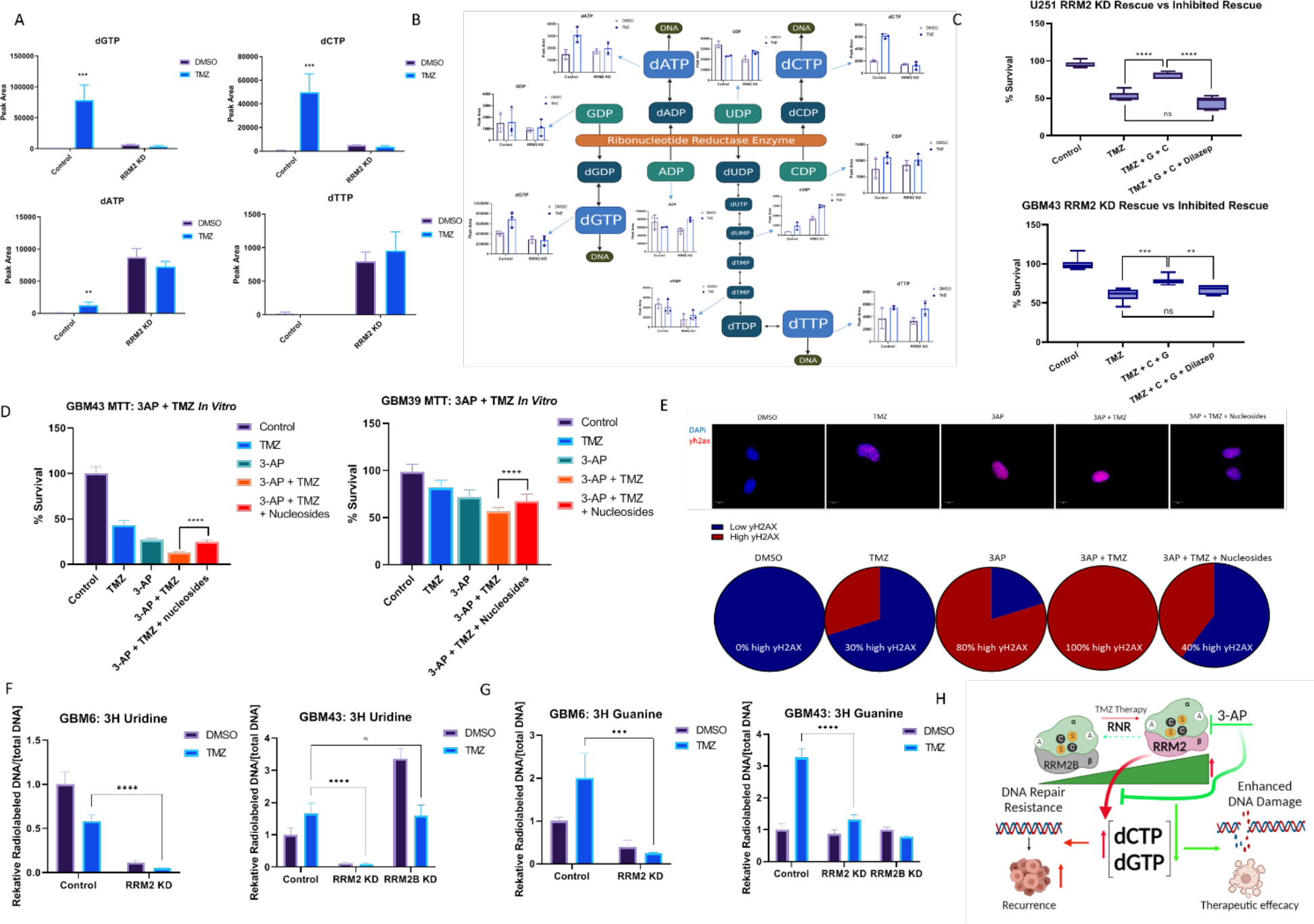
RRM2-mediated production of dCTP and dGTP is necessary for adaptation to TMZ therapy. **(A)** Targeted metabolomics analysis of dNTPs production in control of shRNA-mediated RRM2 KD cells. Metabolomics analysis performed at 72 hours post TMZ (100uM) exposure. Graphed in Prism GraphPad 9.0, using to compare row-means to determine significance or using log-rank tests to determine significance. **(B)** Schematic of the RNR-mediated de novo dNTP synthesis and the corresponding bulk metabolomics analysis graphs of metabolites included in this pathway. Graphed in Prism GraphPad 9.0, using to compare row-means to determine significance or using log-rank tests to determine significance. **(C)** Rescue of TMZ sensitivity of shRNA- mediated RRM2 KD cells when treated with DMSO (100uM), TMZ (100uM), TMZ + deoxycytidine (25uM) and deoxyguanosine (25uM) nucleosides. Cells treated with TMZ + deoxycytidine (25uM) and deoxyguanosine (25uM) nucleosides + Dilazep (5uM) block rescue effect. **(D)** Rescue of 3AP Triapine and TMZ sensitivity of shRNA-mediated RRM2 KD cells when treated with DMSO (100uM), TMZ (100uM), 3-AP Triapine (2uM), 3-AP Triapine + TMZ, or 3-AP Triapine + TMZ + deoxycytidine (25uM) and deoxyguanosine (25uM) nucleosides. Graphed in Prism GraphPad 9.0, using to compare row-means to determine significance or using log-rank tests to determine significance. Validated in GBM43 and GBM39. **(E)** Immunocytochemistry of RRM2 KD cells when treated with DMSO (100uM), TMZ (100uM), 3- AP Triapine (2uM), 3-AP Triapine + TMZ, or 3-AP Triapine + TMZ + deoxycytidine (25uM) and deoxyguanosine (25uM) nucleosides. Nucleus stained in blue (DAPi). yH2AX stained in red (APC). ImageJ Fiji used to quantify yH2AX foci. Analyzed median foci across conditions and plotted in Prism GraphPad 9.0. **(F)** Isotope tracing analysis of 3H Uridine incorporation via de novo synthesis of DNA with or without TMZ (50 uM for 72 hours), comparing Control cells, shRNA-mediated RRM2 KD cells, and shRNA-mediated RRM2B KD cells. Validated in GBM6 and GBM43. Graphed in Prism GraphPad 9.0, using to compare row-means to determine significance or using log-rank tests to determine significance. **(G)** Isotope tracing analysis of 3H Guanine incorporation via de novo synthesis of DNA with or without TMZ (50 uM for 72 hours), comparing Control cells, shRNA-mediated RRM2 KD cells, and shRNA-mediated RRM2B KD cells. Validated in GBM6 and GBM43. Graphed in Prism GraphPad 9.0, using to compare row- means to determine significance or using log-rank tests to determine significance. **(H)** Schematic of hypothesis and summary data made through BioRender. *p < 0.05; **p < 0.01; ***p < 0.001; ****p < 0.0001; ns, not significant.

We next wanted to determine whether nucleosides could be exogenously supplemented to RRM2- knockdown cells to supply necessary deoxynucleotides. An MTT rescue experiment was performed in which U251, GBM43, GBM6, and GBM39 RRM2-knockdowns were supplemented with deoxycytidine and deoxyguanosine, dephosphorylated forms of dCTP and dGTP respectively, in cell media during TMZ treatment. This experiment revealed that exogenous nucleosides could rescue the TMZ susceptible phenotype of RRM2-knockdowns and led to an average 30% rescue in cell survival (*p<0.001*) (Figure 7C, Figure S6b). It is known that nucleoside transporters contribute to intracellular nucleoside homeostasis, and often when nucleoside transporters are blocked, the rescue effect of exogenous nucleosides is reduced^28^. To verify that the supplemental nucleosides were responsible for increased cell survival of RRM2-knockdowns, in the same MTT, we treated a group cells that had received TMZ and supplemental deoxycytidine and deoxyguanosine, with Dilazep, a known nucleoside transport inhibitor^29^. The result of this cell population that had received specific supplemental nucleosides, TMZ, and Dilazep, which confirmed that with nucleoside transport is blocked, cells no longer exhibited the rescue phenotype (*p<0.01*) (Figure 7C, Figure S6c). Finally, to ensure the phenotypic rescue of RRM2-knockdown cells was mediated by deoxycytidine and deoxyguanosine specifically, deoxyadenosine and thymine were also supplemented to cell media and the same MTT experiment was performed. The addition of these nucleosides does not rescue RRM2-knockdown cells from their TMZ- susceptibility (*p<0.001*) (Figure S6d).

We next wanted to determine whether supplemental deoxycytidine and deoxyguanosine could rescue GBM cell lines from the phenotypic effects that 3-AP in combination with TMZ have been shown to have both *in vitro* and *in vivo.* Performing another MTT assay, we found that compared to cells treated with 3-AP and TMZ, cells treated with 3-AP and TMZ and a supplemental dose of nucleosides could rescue the GBM from drug-induced death. (Figure 7D). To further support the rescue effect that supplemental nucleosides induce, we used immunocytochemistry to stain for yH2AX in cells treated with TMZ alone, 3-AP alone, a combination of 3-AP and TMZ, a combination of 3-AP, TMZ, and supplemental nucleosides, or vehicle control DMSO. Using ImageJ to quantify yH2AX foci count per cell, per condition, we can confirm our previous data in which the addition of deoxycytidine and deoxyguanosine promotes less yH2AX, indicating more efficient DNA damage repair and less overall cellular death (Figure 7E).

The significance of deoxycytidine and deoxyguanosine revealed first in our targeted metabolomics analysis and confirmed through bulk metabolomics and several rescue experiments, was further explored through the analysis of nucleobase transport measured through radioactive flux assays to measure 3H-guanine and 3H uridine incorporation into DNA. Using established RRM2- knockdowns, 3H Uridine and 3H Guanine DNA was traced through treatment conditions. Both Uridine and Guanine, were incorporated into RRM2 KD cells significantly less than controls across conditions (*p<0.001*) (Figure 7F). Contrarily, when the same experiment was performed using established RRM2B-knockdowns, we found that the incorporation of Uridine and Guanine once measured was similar to control cells (Figure 7G). Not only does this support the mediation of nucleoside transporters in supplying cells with deoxycytidine and deoxyguanosine, it also sheds light on idea that without RRM2, cells are unable to incorporate Guanine and Uridine into DNA in to the same efficiency as control.

Taken together, our studies have revealed a novel approach to targeting the RNR enzyme and identified the corresponding molecular mechanism underlying the success of this approach (Figure 7H). As previously discussed, stable nucleotide pools are essential for proper DNA repair and genomic stability. RRM2 has been shown to drive the homeostatic nature of nucleotide pools. Cells that have undergone RRM2-knockdown or drug-induced RRM2 inhibition, are unable to produce dCTP and dGTP at the same rates as control cells, therefore throwing the entire nucleotide pool out of homeostasis. RRM2 inhibition not only reduces DNA repair capacity and consequent TMZ-resistance, it also enhances DNA damage, leading to increased therapeutic efficacy.

## DISCUSSION

Imagine attending a football game in which you are only allowed to see the kickoff and final score. The entire game – where fumbles are made, passes intercepted, and calls overturned – would be a complete mystery. Our previous understanding of GBM was predominately limited to tissue sample collected during the initial surgery and tissue collected during post-recurrent surgery – the kickoff and final score, respectively. Cellular adaptation during TMZ therapy has been shown to play a significant role in the devastating and fatal recurrence of GBM^7, 8^. Evolutionary and selection pressure reminds us that the most fit cancer cells will survive chemotherapy, progressing and growing into the dominant population. The cellular and molecular transformations that arise between primary and recurrent GBM tumors remains a critical gap in GBM research and therapeutic discovery.

In this study, we bridge the gap between primary and recurrent GBM using our novel single-cell sequencing approach. Our sequencing analysis illustrated the unique nature of cells present during therapy, compared to primary and recurrent GBM. Although the idea of cellular differentiation between treatment conditions is somewhat expected, the results of our screen illustrate the dramatic contrast between during therapy GBM and primary/recurrent GBM. Gene Set Enrichment Analysis allowed us to explore the functional differences between these populations as well, in which we see, for example, distinct clusters of DNA repair and chemoresistance genes highly expressed during therapy while expressed at low levels in primary and recurrent GBM. We found this to be quite significant because targeting clusters of DNA repair and chemoresistance genes upregulated during therapy in our sequencing model, has great therapeutic potential in GBM. Uncovering the similarities and differences between cancer cell populations’ expression during therapy compared to recurrent GBM, sheds light on the molecular mechanisms underlying chemoresistance.

We then specifically compared the populations of cells during therapy and in recurrent GBM using the same GSEA techniques and further pathway analysis. Our screen allowed us to parse out the differences between GBM tumors exposed to a limited or a full duration of chemotherapy. The most striking difference in this analysis was revealed through the enriched pathways present during therapy versus in recurrent GBM. And with this, not only the function of the enriched pathways, but the quantity and heterogeneity of enriched pathways as well. For example, the pathway analysis of oncogenic gene sets enriched during therapy revealed that the majority of cells were recruited to respond to ‘Chemicals’, ‘Stress,’ or ‘Chemotaxis’. In contrast, the oncogenic gene sets enriched in recurrent GBM revealed pathways associated with everything from the regulation of ‘Cell Cycle,’ ‘Cell Proliferation,’ and ‘Cellular Organization,’ to the ‘Oxidation-Reduction,’ ‘Steroid Metabolic,’ and ‘Organophosphate Biosynthetic’ processes. Equally as important, the networks in recurrent GBM consisted of far more major pathways than networks elevated during therapy. It is evident that the specific pathways and networks of genes that are uniquely enriched during therapy must be further investigated to progress GBM research and discover novel therapeutic approaches.

To further investigate the functional role of the genes enriched during therapy, we preformed pathway analysis, in which we found one of the top hits to be pathways involved in ‘Metabolic Processes.’ One of the hallmarks of cancer in general is metabolic adaptation and GBM cells are no exception. Not only does metabolic transformation contribute cancer cell proliferation, but it also plays a significant role in chemoresistance. Targeting metabolic adaptation has become an increasingly attractive method of cancer therapy^30^. Due to this significance, we wanted to further explore the genes in our sequencing data that were driving metabolic processes during therapy. It was at this point that we discovered the unique signature that the RNR family of genes showed in our data; while RRM1 and RRM2B remained relatively constant across conditions, RRM2 expression was significantly elevated during therapy. This family of genes was of particular interest to us due to its importance in proper DNA synthesis and genomic stability. For the RNR enzyme to form, RRM1 must bind to either RRM2 or RRM2B. Although the RNR enzyme itself, essential for dNTP metabolism, has been extensively studied, the specific subunits have not. From our initial sequencing data, we hypothesized that the enzyme undergoes a preferential switch in subunit binding to favor RRM2 during therapy. Our goal then became to test this hypothesis and elucidate the molecular mechanisms underlying this hypothesis.

Altering the expression of RNR genes provided critical data. We found that RRM2 KD cells create an incredibly TMZ sensitive phenotype in GBM cells, while RRM1 and RRM2B KD cells cause TMZ resistance. We also found that RRM2 OE cells reversed the TMZ sensitivity of KD cells, producing a TMZ resistant phenotype. We next wanted to determine whether this phenomenon was TMZ specific or not. Both CCNU and Ibrutinib did not produce the same phenotypes of our KD cells, indicating the RRM2 KD sensitivity and RRM1/2B KD resistance was in fact TMZ specific. Another key piece of data we presented, was the specificity of dGTP and dCTP in both the RRM2 KD cells reduced production during TMZ therapy, and the ability of these two metabolites to be exogenously added to RRM2 KD cells with TMZ to rescue the sensitive phenotype. Taken together, it is possible there is a connection between specificity of TMZ and corresponding specificity of dGTP and dCTP in our data. It has been shown that the O6- methylguanine (O6-MG) is the critical site of nucleotide damage caused by TMZ therapy, and that GBM cells are able to sense a depletion in their purine nucleotide pools^19^. It is possible that RRM2 mediates *de novo* synthesis of dGTP and dCTP when GBM cells are depleted of nucleotides through TMZ-induced O6-MG damage. However, further investigation must be conducted to determine the foundation of this specificity.

We have demonstrated a novel mechanism of chemoresistance where RRM2-regulated dNTP synthesis promotes adaption to TMZ in GBM. RNR inhibitors, such as Hydroxyurea (HU), have been extensively studied in a clinical setting against many cancers, including GBM, with limited success^31, 32^. Second-generation RNR inhibitors that cross the BBB, such as 3-AP Triapine, could be more effective as it has been measured to be 1000 times more potent than HU in targeting the RNR enzyme^33, 34^. Additionally, 3-AP Triapine has FDA approval and is in several clinical trials, but has never been investigated against GBM^27^. *In vivo* studies of Triapine have revealed its survival benefit in GBM tumor-bearing mice, corresponding to enhanced reactive oxygen species and oxidative DNA lesions in GBM cell lines^35^. In our data, we demonstrate that Triapine sensitizes GBM cells *in vitro* and *in vivo*, leading to significant survival in GBM tumor-bearing mice treated with a combination of Triapine + TMZ.

In conclusion, we present a mechanism of GBM metabolic adaptation that drives chemoresistance, and the corresponding druggable target that can be inhibited to enhance the efficacy of TMZ therapy in clinic.

## ACKNOWLEDGEMENTS

This work was supported by the National Institute of Neurological Disorders and Stroke grant 1R01NS096376, 1R01NS112856, and P50CA221747 SPORE for Translational Approaches to Brain Cancer. The results published here are in part based upon data generated by the TCGA Research Network: https://www.cancer.gov/tcga, and were further analyzed through GlioVis. In addition, these results use data generated by the Human Protein Atlas and GBMSeq (Gephart Lab, www.gbmseq.org). Figures, in part, were generated using BioRender (www.biorender.com).

## AUTHOR CONTRIBUTIONS

Conceptualization: Ella N Perrault, Jack M Shireman, Atique U Ahmed

Methodology: Ella N Perrault, Jack M Shireman, Eunus Ali, Atique U Ahmed

Validation, Formal Analysis, Investigation: Ella N Perrault, Eunus Ali, Isabelle Preddy, Peiyu Lin, Cheol Park, Luke Tomes, Andrew J Zolp, Shreya Budhiraja, Shivani Baisiwala

Resources: Atique U Ahmed, Issam Ben-Sahra, C. David James, Sebastian Pott, Anindita Basu Data Curation & Draft Preparation: Ella N Perrault, Isabelle Preddy

Review & Editing: Ella N Perrault, Isabelle Preddy, Shreya Budhiraja, Atique U Ahmed Supervision, Project Administration, Funding Acquisition: Atique U Ahmed

---

**Figure S1:**
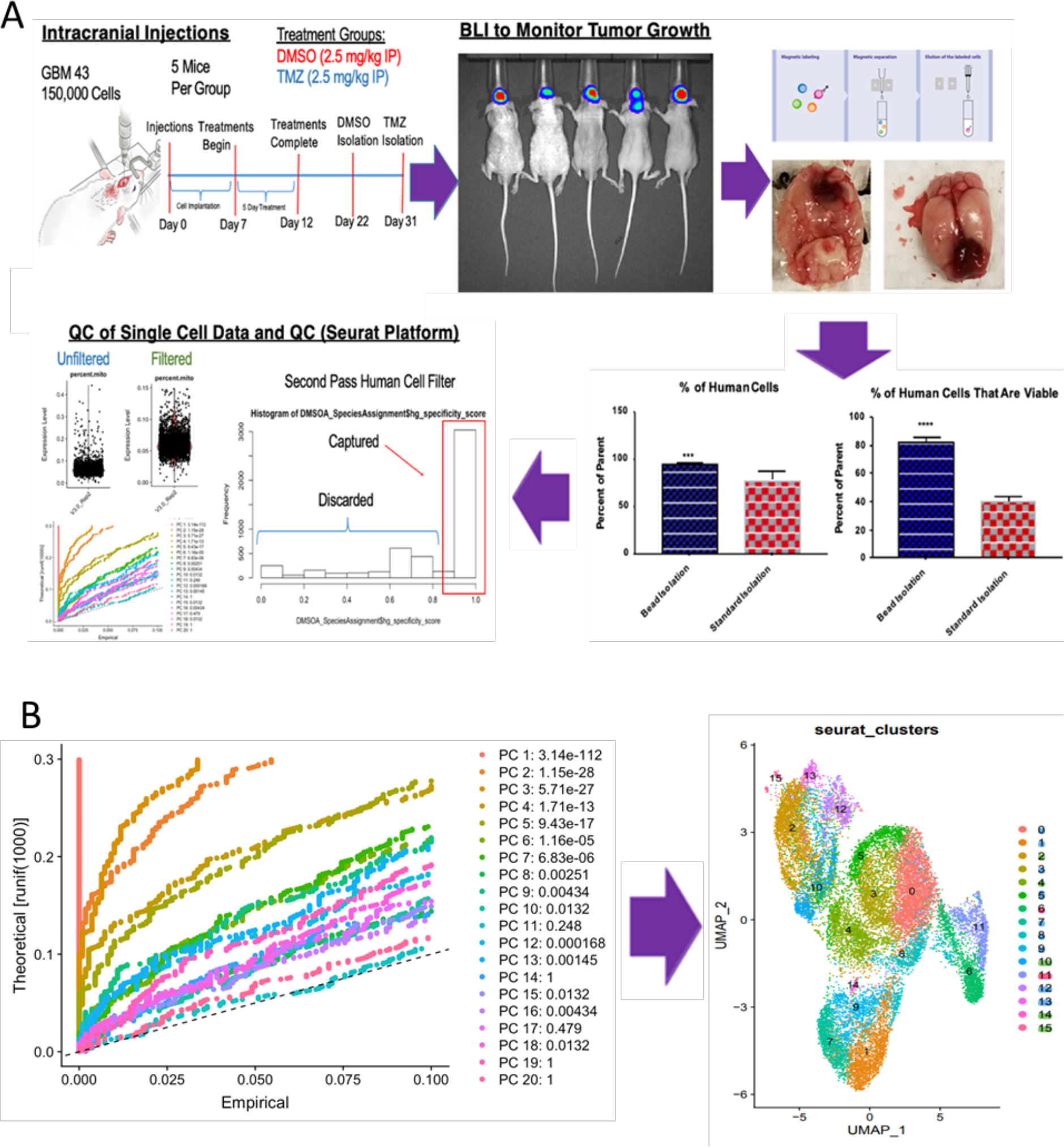
Single-Cell RNA Sequencing In Vivo Pipeline. (A) Schematic of scRNA-sequencing in vivo pipeline performed using droplet-based sequencing of human GBM43 cells. (B) Principal Component Analysis and Seurat Analysis clustering of our scRNAseq data.

**Figure S2:**
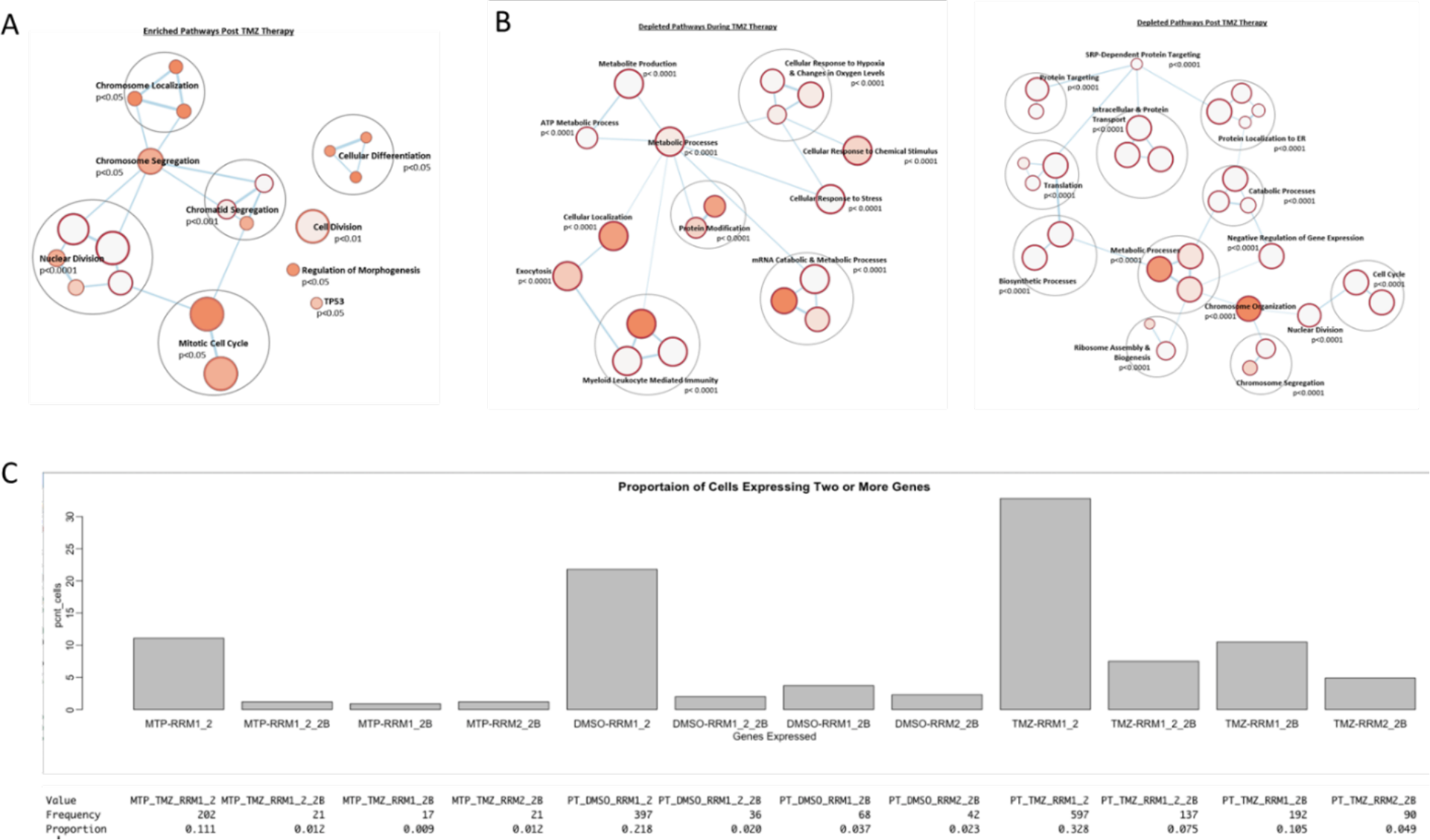
Single-Cell RNA Sequencing Pathway Analysis and RNR gene expression patterns. (A-B) Pathway analysis of genes enriched or depleted in our scRNAseq data. Analyzed through gProfiler and Cytoscape. (C) scRNAseq analysis of the proportion of genes expressing a certain combination of RRM1, RRM2, and RRM2B, in each treatment condition of the screen.

**Figure S3:**
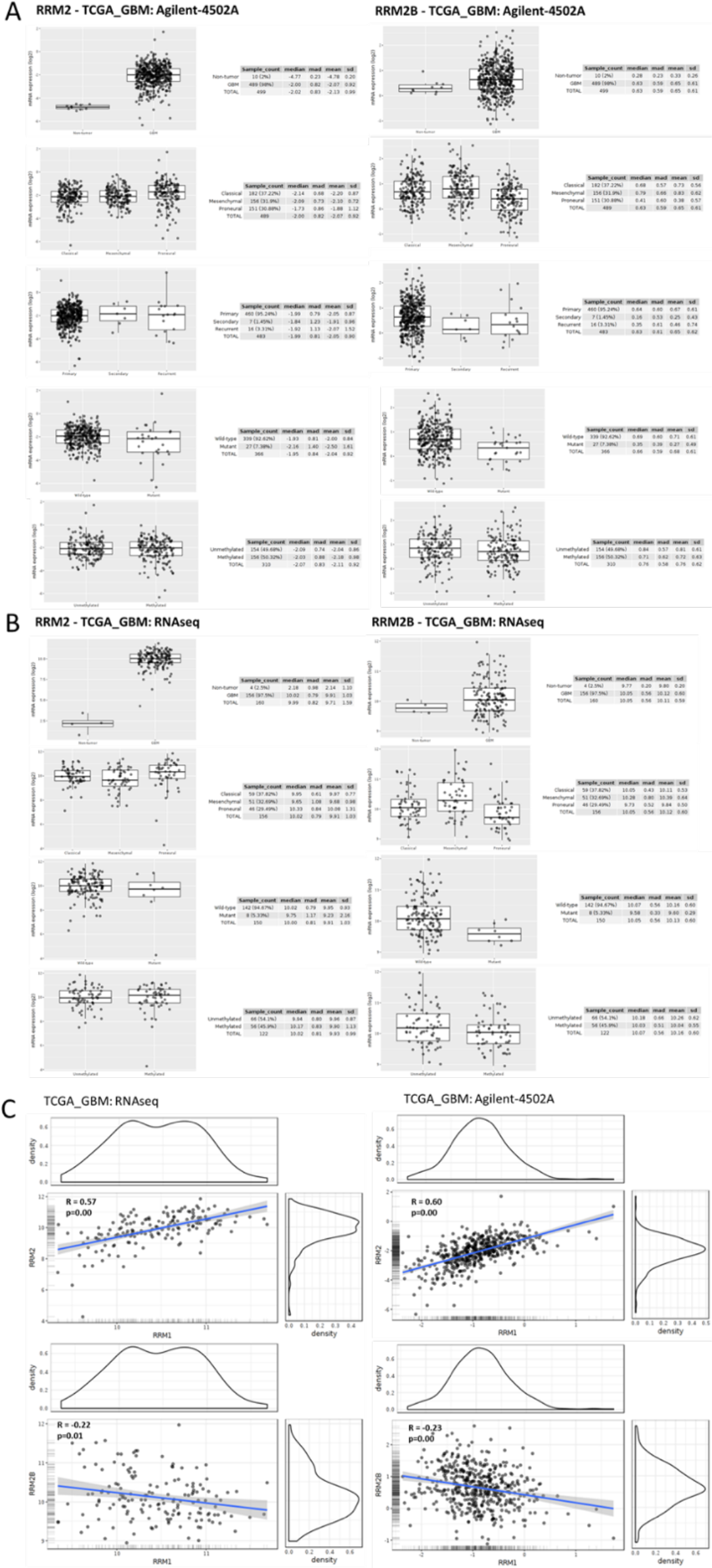
RNR gene expression in patient datasets. **(A**) Representative GlioVis plots of RRM2 (left) and RRM2B (right) gene expression in different conditions (i.e. GBM subtype, IDH mutant status, etc.) from TCGA_GBM: Agilent_4502A database. **(B**) Representative GlioVis plots of RRM2 (left) and RRM2B (right) gene expression in different conditions (i.e. GBM subtype, IDH mutant status, etc.) from TCGA_GBM: RNAseq database. **(C)** Representative correlation plot of RRM1-RRM2 (top) or RRM1-RRM2B (bottom) correlations in TCGA_GBM: RNAseq (left) or TCGA_GBM: Agilent-4502A (right) databases on GlioVis.

**Figure S4:**
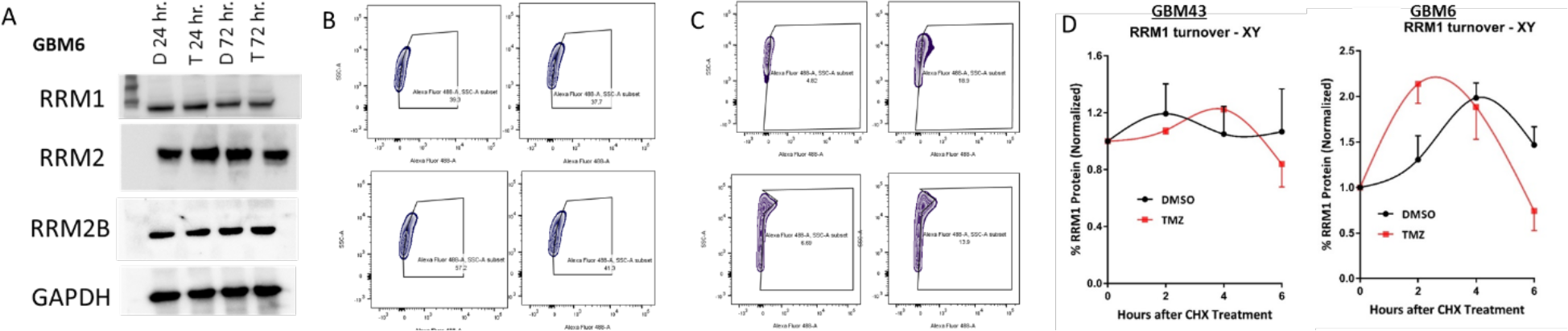
RNR gene expression during TMZ therapy. (A) Western-blot analysis of GBM6 cells treated for 24 or 72 hours of DMSO (50uM) or TMZ (50uM) and stained for genes of interest. (B) Representative FACS gating corresponding to Figure 5D. Analyzed with FlowJo. (C) Representative FACS gating corresponding to Figure 5E. Analyzed with FlowJo. (D) Chase Assay graphs of RRM1 and GAPDH (control) corresponding to Figure 5G. Graphed in Prism GraphPad 9.0.

**Figure S5:**
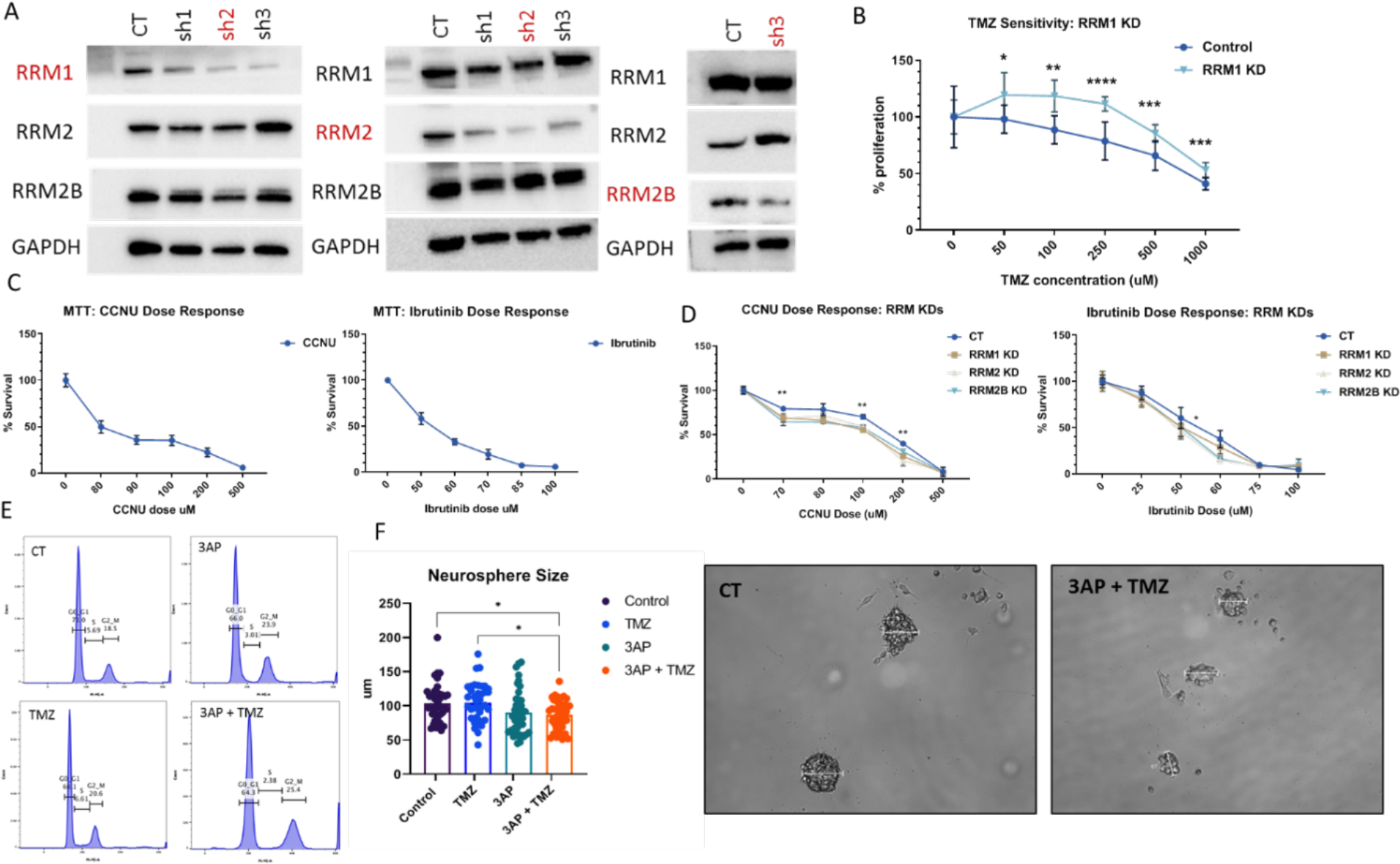
RNR KD drug-induced phenotype is TMZ specific; 3-AP Triapine inhibits neurosphere size. **(A)** Western blot analysis of all shRNA-mediated KDs of RRM1, RRM2, and RRM2B. Red highlights are the most efficient shRNA-mediated knockdowns chosen. **(B)** MTT cell viability assay performed on shRNA-mediated RRM1 KD cells in TMZ dose response (0uM, 50uM, 100uM, 250uM, 500uM, 1000uM). Graphed in Prism GraphPad 9.0, using to compare row-means to determine significance or using log-rank tests to determine significance. **(C)** MTT cell viability assay to identify IC50 of drugs of interest after 48 hours. Cells treated with CCNU (0uM, 80 uM, 90uM, 100uM, 200uM, 500uM) and cells treated with Ibrutinib (0uM, 50uM, 60uM, 70uM, 85uM, 100uM). Graphed in Prism GraphPad 9.0. **(D)** MTT cell viability assay of control cells vs shRNA-mediated RRM1 KD, RRM2 KD, and RRM2B KD cells when treated with CCNU (0uM, 70uM, 80uM, 100uM, 200uM, 500uM) for 48 hours. MTT cell viability assay of control cells vs shRNA-mediated RRM1 KD, RRM2 KD, and RRM2B KD cells when treated with Ibrutinib (0uM, 25uM, 50uM, 60uM, 75 uM, 100uM) for 48 hours. Graphed in Prism GraphPad 9.0, using to compare row-means to determine significance or using log-rank tests to determine significance. **(E)** Representative cell-cycle FACS histograms corresponding to Figure 6G. Analyzed with FlowJo. Bar graph of neurosphere sizes from neurosphere assay of GBM43 cells treated with DMSO (50uM), TMZ (50uM), 3-AP Triapine (2uM), or TMZ + 3AP Triapine. Graphed in Prism GraphPad 9.0, using to compare row-means to determine significance or using log-rank tests to determine significance. Representative images of neurospheres with measurement bar to quantify sizes. Imaged on Bright Field objective of Leica Microscope. Neurosphere size analysis corresponds to Figure 6H. *p < 0.05; **p < 0.01; ***p < 0.001; ****p < 0.0001; ns, not significant.

**Figure S6:**
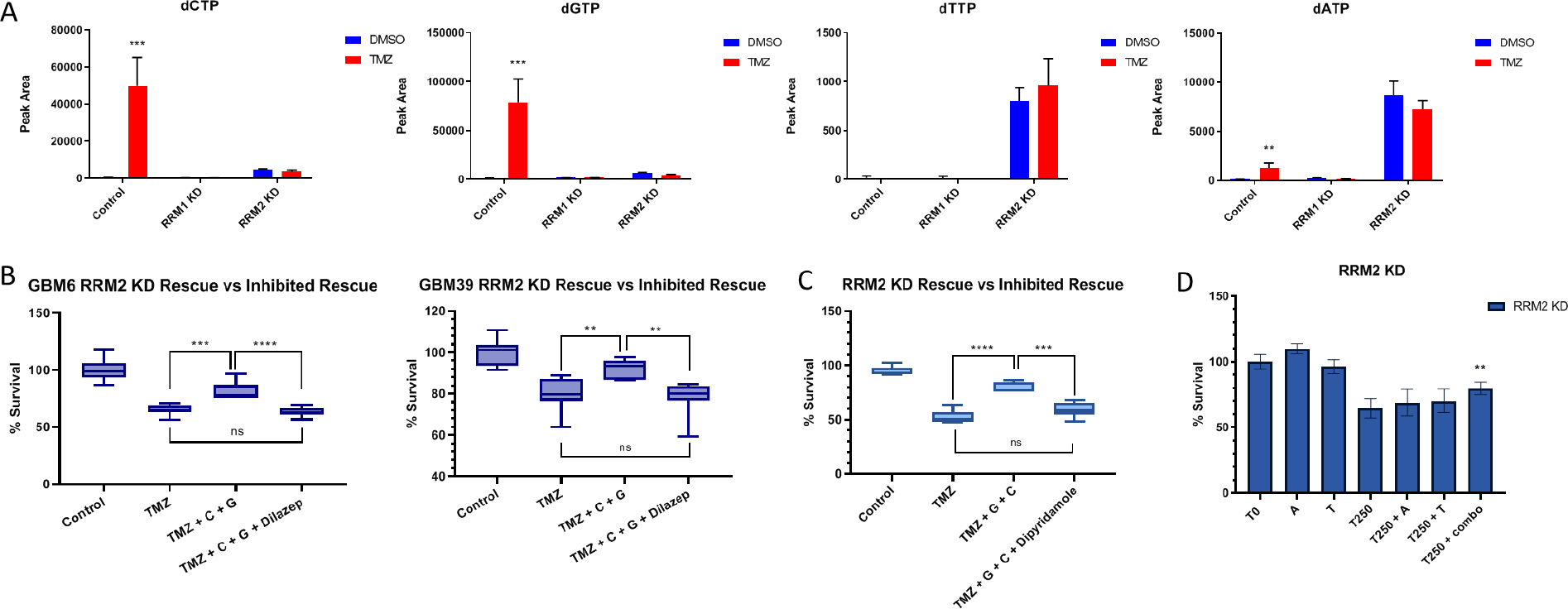
Targeted Metabolomics reveals key metabolites necessary for RRM2 KD cells to combat TMZ. **(A)** Original targeted metabolomics analysis of dNTP production in control cells, shRNA-mediated RRM1-KD cells, and shRNA-mediated RRM2 KD cells when treated with DMSO (100uM) or TMZ (100uM) for 72 hours. **(B)** Figure 7C validated in two more cell lines: GBM6 and GBM39. (C) An MTT cell viability assay of RRM2 KD cells treated with TMZ + deoxycytidine (25uM) and deoxyguanosine (25uM) nucleosides + Dypridamole (5uM) another known nucleoside transport inhibitor^37^, to block rescue effect. Corresponding to Figure 7C. Graphed in Prism GraphPad 9.0, using to compare row-means to determine significance or using log-rank tests to determine significance. **(D)** An MTT viability assay of shRNA-mediated RRM2 KD cells when treated with DMSO (100uM), TMZ (100uM), TMZ + deoxyadenosine (25uM) and thymine (25uM) nucleosides, to show nucleoside specificity in TMZ sensitivity rescue. Graphed in Prism GraphPad 9.0, using to compare row-means to determine significance or using log-rank tests to determine significance. *p < 0.05; **p < 0.01; ***p < 0.001; ****p < 0.0001; ns, not significant.

